# Biomechanical regulation of Ca^2+^ dynamics during muscle stem cell activation

**DOI:** 10.64898/2026.05.19.726396

**Authors:** Kotaro Hirano, Yudai Ishikawa, Norio Motohashi, Yujiro Kobata, Hiroto Watanabe, Mao Sasaki, Takuya Yokoyama, Yuto Yamada, Yuta Aoyagi, Kanako Takakura, Akira Murakami, Masaki Tsuchiya, Yusuke Ono, Keiko Nonomura, Yoshitsugu Aoki, Yuji Hara

## Abstract

Muscle satellite cells (MuSCs) are muscle-resident stem cells that are responsible for myofiber regeneration. Although the importance of calcium ions (Ca^2+^) in muscle physiology has been well established, the mechanism by which Ca^2+^ mobilization governs MuSC function remains poorly understood. In this study, we aimed to systematically characterize Ca^2+^ dynamics in MuSCs and to define the mechanisms regulating these signals during muscle regeneration. By employing modified protocols for mouse MuSC isolation and Ca^2+^ measurement, we observed spontaneous Ca^2+^ fluctuations in MuSCs isolated from regenerating muscle after cardiotoxin-induced myofiber injury. Our detailed analysis using chemical Ca^2+^ indicators and a genetically encoded Ca^2+^ indicator revealed that the frequency and amplitude of Ca^2+^ fluctuations increased significantly during the activated and proliferative stages of MuSCs. This effect was more pronounced in MuSCs isolated from dystrophic and aged mice. Mechanistically, these Ca^2+^ fluctuations were at least partially mediated by mechanosensitive ion channels, including PIEZO1 and TRPM7, which promote MuSC migration. Collectively, our findings demonstrate that Ca^2+^ fluctuations through mechanosensitive ion channels act as a key regulator of MuSC activation during muscle regeneration and may provide new insights into the role of Ca^2+^ influx in muscle biology and the pathogenesis of muscle diseases.

## Introduction

Calcium ions (Ca^2+^) act as universal second messengers; however, their intracellular concentrations are maintained under strict control^1,2^. Under resting conditions, cytoplasmic Ca^2+^ levels are sustained at approximately 100 nM, nearly four orders of magnitude lower than extracellular Ca^2+^ concentrations (approximately 1–2 mM), thereby establishing a steep transmembrane Ca^2+^ gradient. This gradient permits rapid Ca^2+^ influx through plasma membrane ion channels in response to stimuli, coordinating diverse cellular processes, including gene expression, metabolism, cytoskeletal dynamics, and cell fate decisions^3,4^. At the cellular level, Ca^2+^ signaling often manifests as cell-autonomous Ca^2+^ fluctuations under basal conditions in various cell types^5-9^. The magnitude of Ca^2+^ influx can be quantified as the amplitude of these fluctuations, defined as the difference between peak and basal Ca^2+^ levels. Although changes in Ca^2+^ amplitude are tightly associated with the promotion or suppression of cellular processes, the mechanisms and physiological conditions underlying such Ca^2+^ dynamics remain largely unclear. Muscle satellite cells (MuSCs) are resident stem cells of skeletal muscle and are indispensable for muscle regeneration^10^. Upon muscle injury, MuSCs exit quiescence, enter the cell cycle, and proliferate as myoblasts, which subsequently migrate and fuse to repair or regenerate damaged myofibers^11^. MuSCs constitute a rare cell population, accounting for approximately 2–10% of total myonuclei in adult skeletal muscles, which has previously limited their experimental accessibility^12^. Advances in cell isolation techniques, including fluorescence-activated cell sorting (FACS)^13-15^ and magnetic activated cell sorting (MACS)^16^, have enabled the isolation of MuSCs with high yield and purity, thereby allowing detailed molecular and functional characterization. However, accumulating evidence indicates that MuSCs isolated using these *ex vivo* approaches exhibit both transcriptional^17-19^ and morphological alterations^20^ compared with their *in vivo* counterparts, raising important questions regarding how accurately isolated MuSCs reflect their native physiological state.

The role of Ca^2+^ in skeletal muscle has been demonstrated in multiple contexts. For example, Ca^2+^ plays key roles in excitation–contraction coupling during muscle contraction and retraction^1,21^, myofiber membrane repair^22,23^, myoblast differentiation and fusion^24,25^, muscle hypertrophy^26,27^, and muscular dystrophies^28-30^. Although the importance of Ca^2+^ in skeletal muscle has long been recognized^1^, no comprehensive study has examined its role in MuSCs during their activation from quiescence.

We previously reported that spontaneous Ca^2+^ influx is observed in freshly isolated MuSCs^31^; however, whether these events can be observed in activated MuSCs has not been investigated. In this study, we combined MACS and FACS to efficiently measure Ca^2+^ influx in MuSCs directly isolated from mice. Using a fluorescent calcium indicator, we further show that spontaneous Ca^2+^ influx increases in an activation-dependent manner. In addition, we generated genetically modified mice expressing GCaMP6s^32^ restricted to MuSCs, enabling visualization of spontaneous Ca^2+^ influx. We further demonstrate that the mechanosensitive ion channel PIEZO1 is responsible for sensing extracellular stiffness and that spontaneous Ca^2+^ influx in MuSCs is altered by PIEZO1 depletion. Finally, we show that Ca^2+^ influx in MuSCs is essential for early activation signatures and cell cycle entry. Moreover, we demonstrate that Ca^2+^ signaling is required for cell migration and that the mechanosensitive ion channels PIEZO1 and TRPM7 are responsible for MuSC migration. Collectively, this study demonstrates that spontaneous Ca^2+^ activity is essential for MuSC function, with mechanosensitive ion channels serving as upstream regulators.

## Results

### Measurement of cytosolic Ca^2+^ levels in MuSCs during skeletal muscle regeneration

To investigate the role of Ca^2+^ in MuSCs, we developed two methods for Ca^2+^ imaging. First, we used MACS, employing negative selection against CD31, CD45, and Sca-1, followed by positive selection for α7-integrin-positive cells to isolate MuSCs. After MACS, MuSCs were immediately stained with an anti-VCAM-1 antibody to further improve purity, followed by loading with the fluorescent Ca^2+^ indicator Cal-520 to measure cytosolic Ca^2+^ levels, which were then quantified using FACS (Fig. 1A). Second, after isolating MuSCs as indicated (Fig. 1B), we used the ratiometric Ca^2+^ indicator Fura-2 to measure spontaneous Ca^2+^ mobilization using live-cell microscopy (Fig. 1A).

**Figure 1:**
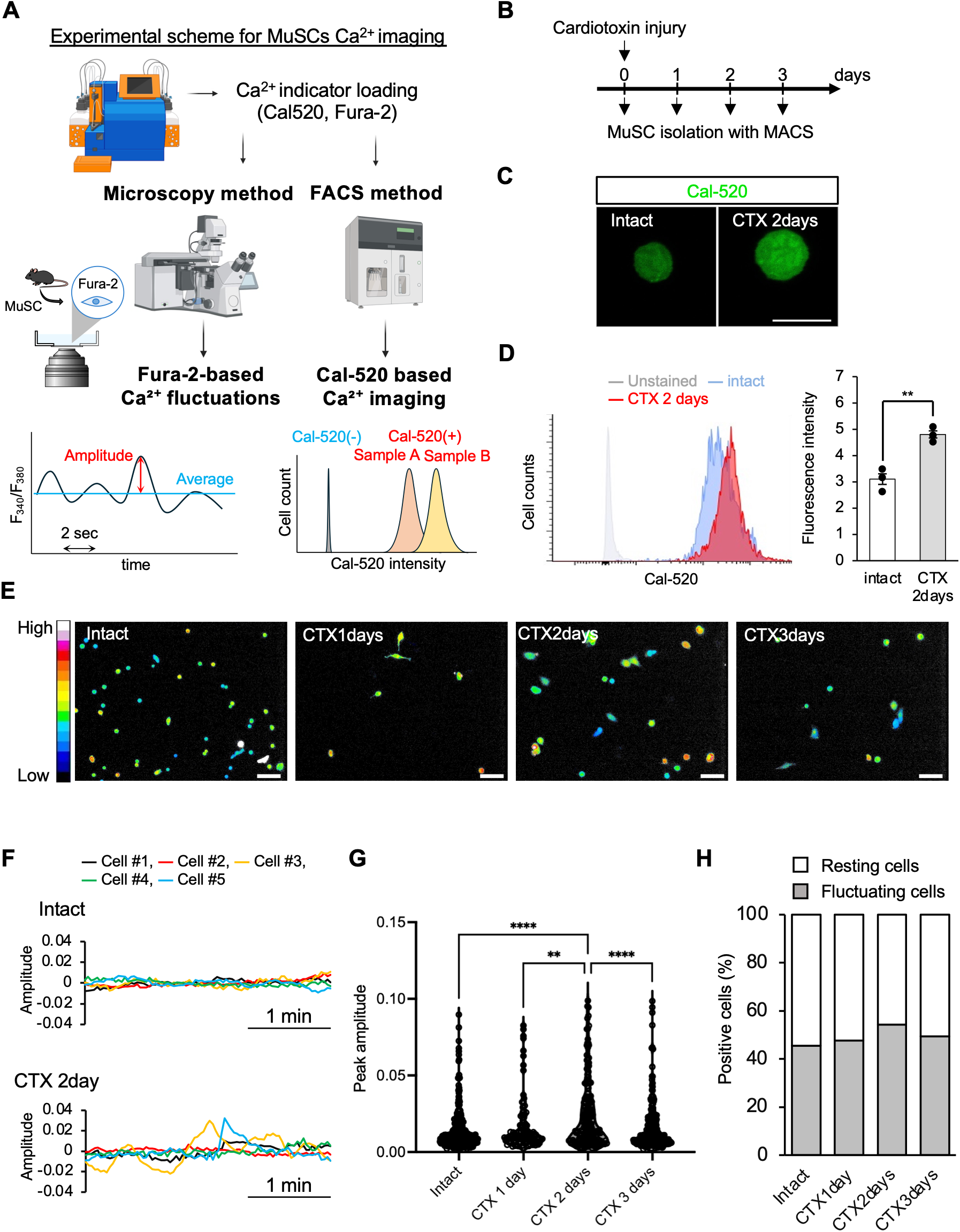
Intracellular Ca^2+^ fluctuations in MuSCs isolated from the injured muscle. **(A)** Experimental scheme for intracellular Ca^2+^ imaging in MuSCs. Upper: Isolation of MuSCs using MACS. Lower Left: Measurement of Fura-2-based Ca^2^+ dynamics. Amplitude is defined as the difference between each point’s F_340_/F_380_ ratio and the mean value. Lower right: Measurement of Cal-520-based Ca^2^+ dynamics. Each mean fluorescence intensities (MFI) are measured from the histogram of flow cytometric data using Cal-520. **(B)** Time course for muscle injury and isolation of MuSCs. **(C and D)** Ca^2+^ measurements in MuSCs using Cal-520, AM. **(C)** Representative fluorescent images of Cal-520-incorporated MuSCs. Left: MuSCs isolated from injured muscle. Right: MuSCs from the injured muscle 2 days after muscle injury. **(D)** Left: Representative histograms of flow cytometric data using Cal-520. Light grey: unstained MuSCs, Blue: MuSCs of intact muscle; and Red: MuSCs of CTX 2 days. Right Quantification of increase in cytosolic Ca^2+^. The data are presented as the means of MFI from N = 3 mice per condition. **(E, F, G and H)** Ca^2+^ measurements in MuSCs using Fura-2, AM. **(E)** Representative F_340_/F_380_ Ratio level images of Fura-2-incorporated MuSCs. Look-up tables (LUTs) indicate signal intensity. **(F)** Representative recording traces of Amplitude from five cells. **(G)** Quantification of the maximum amplitude during the measurement. The data are presented as the means of > 60 cells from N = 3 mice per condition. **(H)** Quantification of Ca^2+^ fluctuation-positive cells exhibiting at least one spontaneous event (> 0.01 in amplitude). The numbers of Ca^2+^ fluctuation-positive cells/total analyzed cells were 131/288, 41/86, 142/262, and 112/227 for intact and 1, 2, and 3 days after CTX injury, respectively. Data were pooled from three independent trials. Data were pooled from three mice. Scale bars: 10 μm in (C) and 50 μm in (E). Fig. 1A was created with BioRender.com. (D) ***: *P* < 0.001. non-paired *t*-test. (G) *: *P* < 0.05, **: *P* < 0.01, ***: *P* < 0.001, ****: *P* < 0.0001. One-way ANOVA followed by Tukey’s test.

Given that MuSCs are highly proliferative, with peak mitotic activity occurring around 2 days after cardiotoxin (CTX)-induced injury, we measured cytosolic Ca^2+^ levels in MuSCs at this time point. MuSCs isolated 2 days after injury exhibited higher Cal-520 fluorescence intensity than that in freshly isolated satellite cells (FISCs), as detected using confocal microscopy (Fig. 1C). Quantitative analysis using FACS confirmed that cytosolic Ca^2+^ levels were significantly elevated in MuSCs at 2 days post-injury (Fig. 1D). These results demonstrate that an increase in cytosolic Ca^2+^ accompanies MuSC activation during skeletal muscle regeneration. While the Cal-520-based analysis was used to assess cytosolic Ca^2+^ levels in isolated MuSCs, the following Fura-2-based live-cell imaging experiments were designed to monitor spontaneous Ca^2+^ dynamics in plated MuSCs under *ex vivo* culture conditions. FISCs and MuSCs isolated at 1, 2, or 3 days post-injury was obtained via MACS, seeded onto glass-bottom dishes, and loaded with the ratiometric Ca^2+^ indicator Fura-2. Ca^2+^ dynamics were recorded using fluorescence microscopy for at least 3 minutes (Fig. 1E). Spontaneous Ca^2+^ fluctuations were clearly detected (Fig. 1F–1H). The amplitude of Ca^2+^ fluctuations in MuSCs increased as muscle regeneration progressed and reached a maximum 2 days after CTX injury (Fig. 1G). In contrast, the proportion of MuSCs exhibiting Ca^2+^ fluctuations did not markedly change following injury (Fig. 1H). These results indicate that the amplitude of Ca^2+^ influx is enhanced during muscle regeneration, whereas spontaneous Ca^2+^ influx events are detected in both FISCs and activated/proliferating MuSCs.

### Spontaneous Ca^2+^ influx is observed in activated and proliferating MuSCs

As shown in Fig. 1, elevated Ca^2+^ mobilization was observed in activated and proliferating MuSCs. To test whether the increase in Ca^2+^ fluctuations observed in injured muscle can be reproduced outside the tissue environment, we cultured MuSCs under defined high-serum conditions that promote MuSC activation and proliferation. MuSCs isolated using MACS were cultured for up to 3 days and then subjected to Ca^2+^ measurements using Fura-2 (Fig. 2A). Consistent with the data in Fig. 1, the amplitude of spontaneous Ca^2+^ influx was significantly increased in MuSCs cultured for 1, 2, and 3 days compared with that in FISCs (Fig. 2B–2D). Notably, the proportion of MuSCs exhibiting Ca^2+^ fluctuations increased progressively with longer culture duration (Fig. 2E). However, this increase in the number of fluctuating cells was not observed in MuSCs directly isolated following muscle injury (Fig. 1E–1H). Taken together, these findings indicate that enhanced Ca^2+^ influx is a characteristic feature of MuSC activation and proliferation, particularly under *ex vivo* culture conditions.

**Figure 2:**
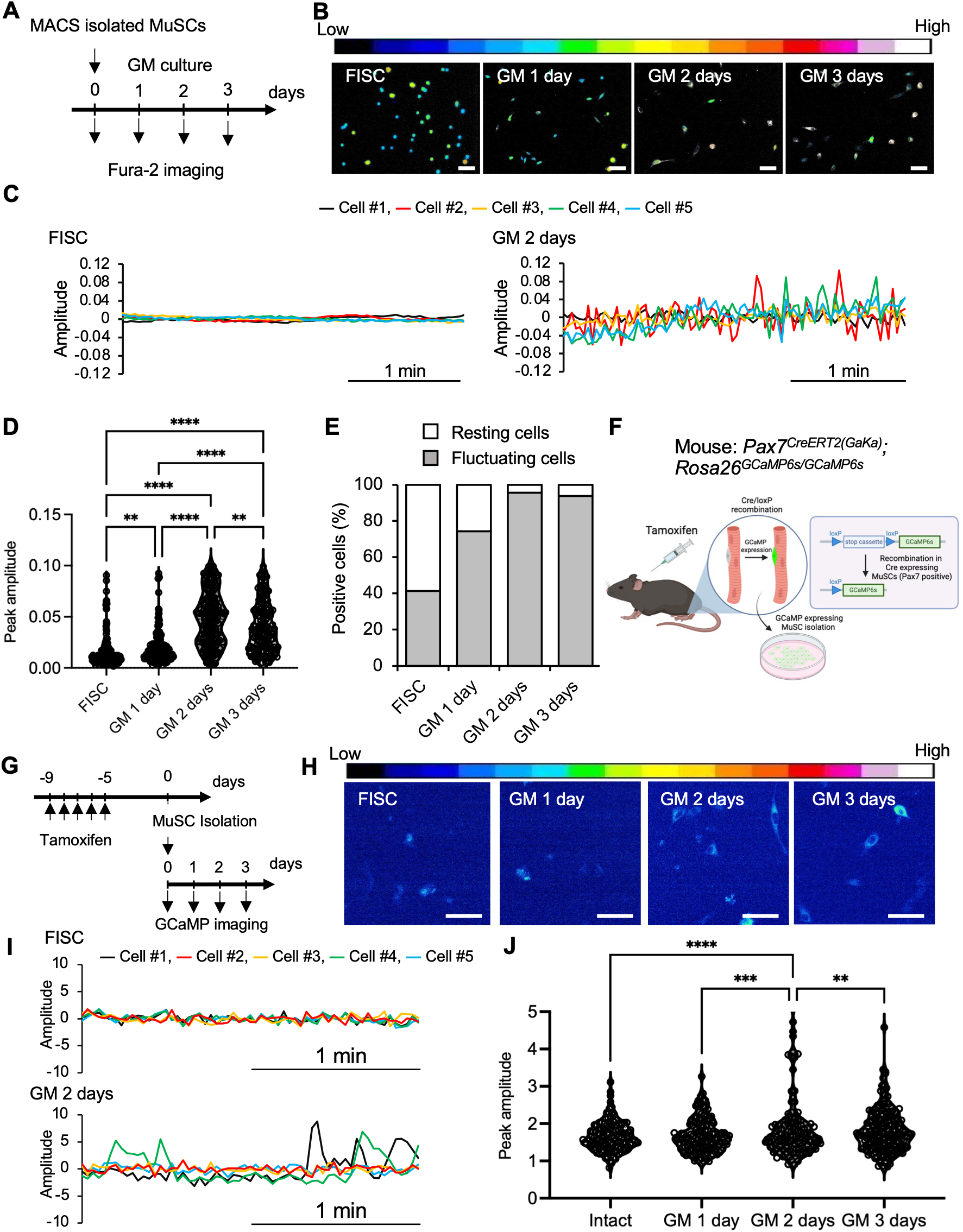
Intracellular Ca^2+^ fluctuations in isolated and cultured MuSCs. **(A)** Time course for isolation and culture of MuSCs under growth medium (GM). **(B, C, D and E)** Ca^2+^ measurements in MuSCs using Fura-2, AM in cultured MuSCs. **(B)** Representative F_340_/F_380_ ratio level images of Fura-2-incorporated MuSCs. LUTs indicate signal intensity. **(C)** Representative recording traces of Amplitude from five cells. **(D)** Quantification of the maximum amplitude during the measurement in FISCs and MuSCs cultured for 1, 2 or 3 days. **(E)** Quantification of Ca^2+^ fluctuation-positive cells exhibiting at least one spontaneous event (> 0.01 in amplitude) under each condition. The numbers of Ca^2+^ fluctuation-positive cells/total analyzed cells were 102/246, 143/192, 172/180, and 139/148 for FISCs and GM 1, 2, and 3 days, respectively. Data were pooled from three independent trials. Data were pooled from three mice. **(F, G, H, I and J)** Intracellular Ca^2+^ fluctuations in MuSCs evaluated by endogenously expressed GCaMP6s in cultured MuSCs. **(F)** Experimental scheme for intracellular Ca^2+^ imaging with GCaMP6s in MuSCs. **(G)** Time course for isolation, culture and GCaMP imaging of MuSCs. **(H)** Representative fluorescence images of GCaMP6s in MuSCs. **(I)** Representative recording traces of Amplitude from five cells. LUTs indicate signal intensity. **(J)** Quantification of the maximum amplitude during the measurement in FISCs and MuSCs cultured for 1, 2 or 3 days. The data are presented as the means of > 100 cells from N = 3 mice per condition. Scale bars: 50 μm in (B) and (H). Fig. 2F was created with BioRender.com. (D) and (I) *: *P* < 0.05, **: *P* < 0.01, ***: *P* < 0.001, ****: *P* < 0.0001. One-way ANOVA followed by Tukey’s test.

### A genetically encoded Ca^2+^ indicator, GCaMP6s, detects spontaneous Ca^2+^ influx in MuSCs

In our previous experiments, Ca^2+^ fluctuations were detected using chemical probes; however, Ca^2+^ mobilization could potentially be influenced by probe-associated cytotoxicity and variability in dye-loading efficiency. To exclude these possibilities, we generated a mouse model harboring a gene cassette encoding the genetically encoded Ca^2+^ indicator GCaMP6s at the Rosa26 locus^32^. To achieve MuSC-specific expression of GCaMP6s, mice carrying *Pax7IRES-CreERT2*^33^ were crossed with *Rosa26GCaMP6s* reporter mice to generate *Pax7IRES-CreERT2; Rosa26GCaMP6s* mice (Fig. 2F). Following five consecutive days of intraperitoneal tamoxifen administration to induce GCaMP6s expression in MuSCs, MuSCs were isolated using MACS (Fig. 2G). Spontaneous Ca^2^+ fluctuations were assessed by monitoring GCaMP fluorescence in FISCs and MuSCs cultured for 1, 2, or 3 days (Fig. 2H). GCaMP-based imaging detected spontaneous Ca^2+^ fluctuations in MuSCs at all the examined time points. Quantitative analysis of GCaMP fluorescence traces acquired over 2 min showed that both the amplitude of Ca^2+^ influx and the proportion of fluctuating cells increased, with a peak observed at 2 days in culture (Fig. 2I and 2J). These findings demonstrate that spontaneous Ca^2+^ influx in MuSCs can be reliably detected using a genetically encoded Ca^2+^ indicator, independently of chemical Ca^2+^ dyes.

### Spontaneous Ca^2+^ fluctuations are observed under diverse physiological conditions in MuSCs

As we demonstrated that Ca^2+^ influx occurs in activated and proliferating MuSCs during muscle regeneration (Fig. 1 and 2), we next examined whether this phenomenon is also observed under physiological or pathological conditions other than regeneration. Skeletal muscle fibers exhibit substantial heterogeneity in contractile properties and metabolic profiles and are broadly classified into fast- and slow-twitch myofibers. Fast-twitch myofibers are characterized by rapid contraction kinetics and predominantly glycolytic metabolism, and they express myosin heavy chain (MyHC) types IIa, IIx, and IIb (encoded by Myh2, Myh1, and Myh4, respectively)^34^. In contrast, slow-twitch myofibers display slower contraction kinetics and oxidative metabolism and express MyHC type I (β), encoded by Myh7^34^. Recent studies have shown that MuSCs retain positional memory^35,36^ and that MuSCs isolated from fast- or slow-twitch myofibers exhibit a bias toward differentiation programs reflective of their fiber-type origin^37^. Based on this, we isolated MuSCs from the extensor digitorum longus and soleus muscles, in which fast- and slow-twitch myofibers are predominantly present, respectively (Fig. 3A). Isolated MuSCs were loaded with the ratiometric Ca^2+^ indicator Fura-2 (Fig. 3B), and spontaneous Ca^2+^ dynamics were measured using fluorescence microscopy. MuSCs derived from fast-twitch myofibers exhibited a significantly higher amplitude of Ca^2+^ influx than those derived from slow-twitch myofibers (Fig. 3C). In addition, a greater proportion of fluctuating cells was observed in MuSCs originating from fast-twitch myofibers (Fig. 3D).

**Figure 3:**
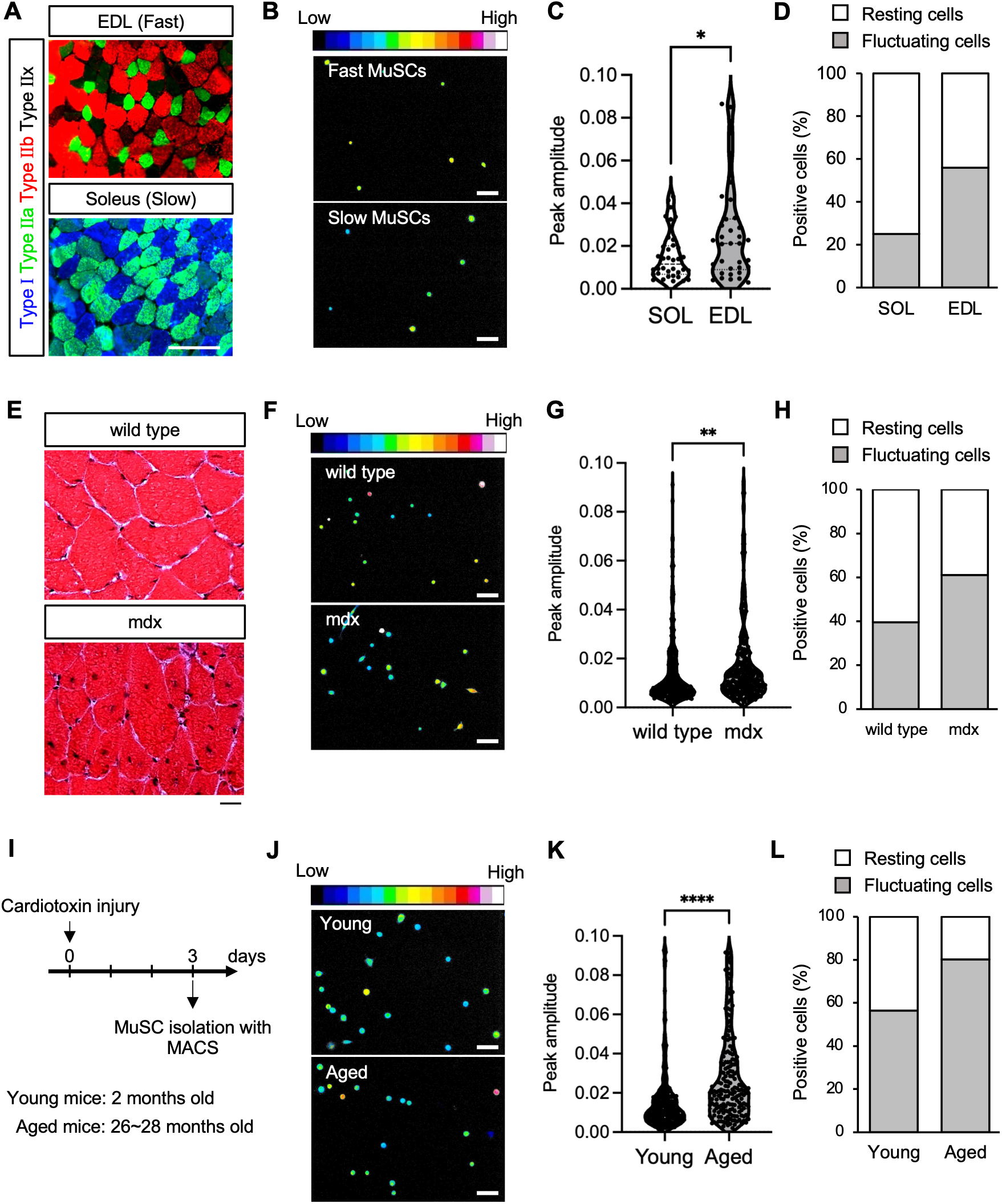
Intracellular Ca^2+^ fluctuations observed in MuSCs under various conditions. **(A)** Immunofluorescence of Myosin heavy chain Type I (blue), IIa (red), IIb (green) muscle fiber staining of cross sections from EDL muscle samples classified as fast-twitch muscle and Soleus classified as slow-twitch muscle. **(B, C and D)** Ca^2+^ measurements in MuSCs using Fura-2, AM in MuSCs isolated from EDL or Soleus. **(B)** Representative F340/F380 ratio level images of Fura-2-incorporated MuSCs. Scale bars: 50 μm. LUTs indicate signal intensity. **(C)** Quantification of the maximum amplitude during the measurement in MuSCs. **(D)** Quantification of Ca^2+^ fluctuation-positive cells exhibiting at least one spontaneous event (> 0.01 in amplitude). The numbers of Ca^2+^ fluctuation-positive cells/total analyzed cells were 9/36 and 19/34 for SOL and EDL, respectively. Data were pooled from six mice. **(F, G and H)** Ca^2+^ measurements in MuSCs using Fura-2, AM in MuSCs isolated from wild type or mdx muscle. **(F)** Representative F340/F380 ratio level images of Fura-2-incorporated MuSCs. LUTs indicate signal intensity. **(G)** Quantification of the maximum amplitude during the measurement in MuSCs. **(H)** Quantification of Ca^2+^ fluctuation-positive cells exhibiting at least one spontaneous event (> 0.01 in amplitude). The numbers of Ca^2+^ fluctuation-positive cells/total analyzed cells were 128/323 and 185/303 for WT and mdx MuSCs, respectively. Data were pooled from three independent trials. **(I, J, K and L)** Ca^2+^ measurements in MuSCs using Fura-2, AM in MuSCs isolated from the injured muscle of young or aged mouse. **(I)** Time course for muscle injury and isolation of MuSCs. **(J)** Representative F340/F380 ratio level images of Fura-2-incorporated MuSCs. LUTs indicate signal intensity. **(K)** Quantification of the maximum amplitude during the measurement in MuSCs. **(L)** Quantification of Ca^2+^ fluctuation-positive cells exhibiting at least one spontaneous event (> 0.01 in amplitude). The numbers of Ca^2+^ fluctuation-positive cells/total analyzed cells were 173/306 and 151/188 for young and aged MuSCs at 3 days after CTX injury, respectively. Data were pooled from three independent trials. (C) (G) (J) *: *P* < 0.05, **: *P* < 0.01, ****: *P* < 0.0001. Unpaired *t*-test.

Next, we examined spontaneous Ca^2+^ influx in a muscle disease model. We evaluated a mouse model for Duchenne muscular dystrophy (*Dmd*^*mdx*^; hereafter referred to as mdx), in which robust and continuous muscle regeneration occurs due to a mutation in the Dmd gene. Mdx mice lack dystrophin and display centrally localized nuclei within myofibers, a hallmark of regenerating muscle fibers (Fig. 3E). MuSCs isolated from mdx mice were loaded with Fura-2 and subjected to Ca^2+^ measurement using fluorescence microscopy (Fig. 3F). Compared to wild-type MuSCs, mdx MuSCs exhibited a significantly higher amplitude of Ca^2+^ influx (Fig. 3G). In addition, the proportion of MuSCs displaying spontaneous Ca^2+^ fluctuations increased in mdx mice (Fig. 3H).

MuSCs are known to exhibit age-associated declines in self-renewal, activation, and proliferative capacity, which collectively contribute to impaired skeletal muscle regeneration^38^. Therefore, we investigated the Ca^2+^ influx capacity of MuSCs isolated from aged mice. To assess Ca^2+^ dynamics in this context, we induced muscle injury using CTX in the hindlimbs of young (2-month-old) and aged (24-month-old) mice (Fig. 3I). MuSCs were isolated 3 days after injury, and spontaneous Ca^2+^ dynamics were measured using Fura-2 (Fig. 3J). Notably, aged MuSCs displayed elevated cytosolic Ca^2+^ levels (Fig. 3K) as well as an increased proportion of cells exhibiting Ca^2+^ fluctuations compared to that with young MuSCs (Fig. 3L). These results indicate that Ca^2+^ fluctuations in MuSCs are influenced by muscle state, including fiber type, muscular dystrophy, and aging.

### Intracellular Ca^2+^ is essential for MuSC activation

In our previous study, we demonstrated that chelation of intracellular Ca^2+^ using BAPTA-AM suppresses MuSC proliferation^31^. In addition, other studies have shown that myoblast fusion is mediated by intracellular Ca^2+^ mobilization^25^. Despite these findings, whether Ca^2+^ influx is required for MuSC activation has not been fully explored. As we recently reported that Mg^2^+ influx is essential for the early activation of MuSCs^39^, we next examined the contribution of Ca^2+^ signaling to early MuSC activation. MuSCs were cultured on isolated myofibers under Ca^2+^-free conditions. MuSCs are known to rapidly induce expression of immediate early genes, including AP-1 family members such as Fos, shortly after myofiber isolation and culture^19,20^. After 2 h of culture under Ca^2+^-free conditions (Fig. 4A), we observed a marked reduction in the number of FOS-positive MuSCs, suggesting that Ca^2+^ influx pathways play a role in the induction of early activation gene signatures (Fig. 4B). MuSCs exhibit characteristic quiescent projections, neurite-like extensions that act as extracellular niche sensors^20,40^. These projections rapidly retract, which is one of the initial events required for subsequent MuSC activation. To assess whether intracellular Ca^2+^ signaling contributes to this morphological transition, myofibers were isolated and cultured in the presence of BAPTA-AM. Notably, quiescent projections were preserved 1 h after culture under Ca^2+^ chelation conditions, whereas projections were lost under control conditions (Fig. 4C). These observations suggest that Ca^2+^ signaling contributes to the early morphological changes associated with MuSC activation.

**Figure 4:**
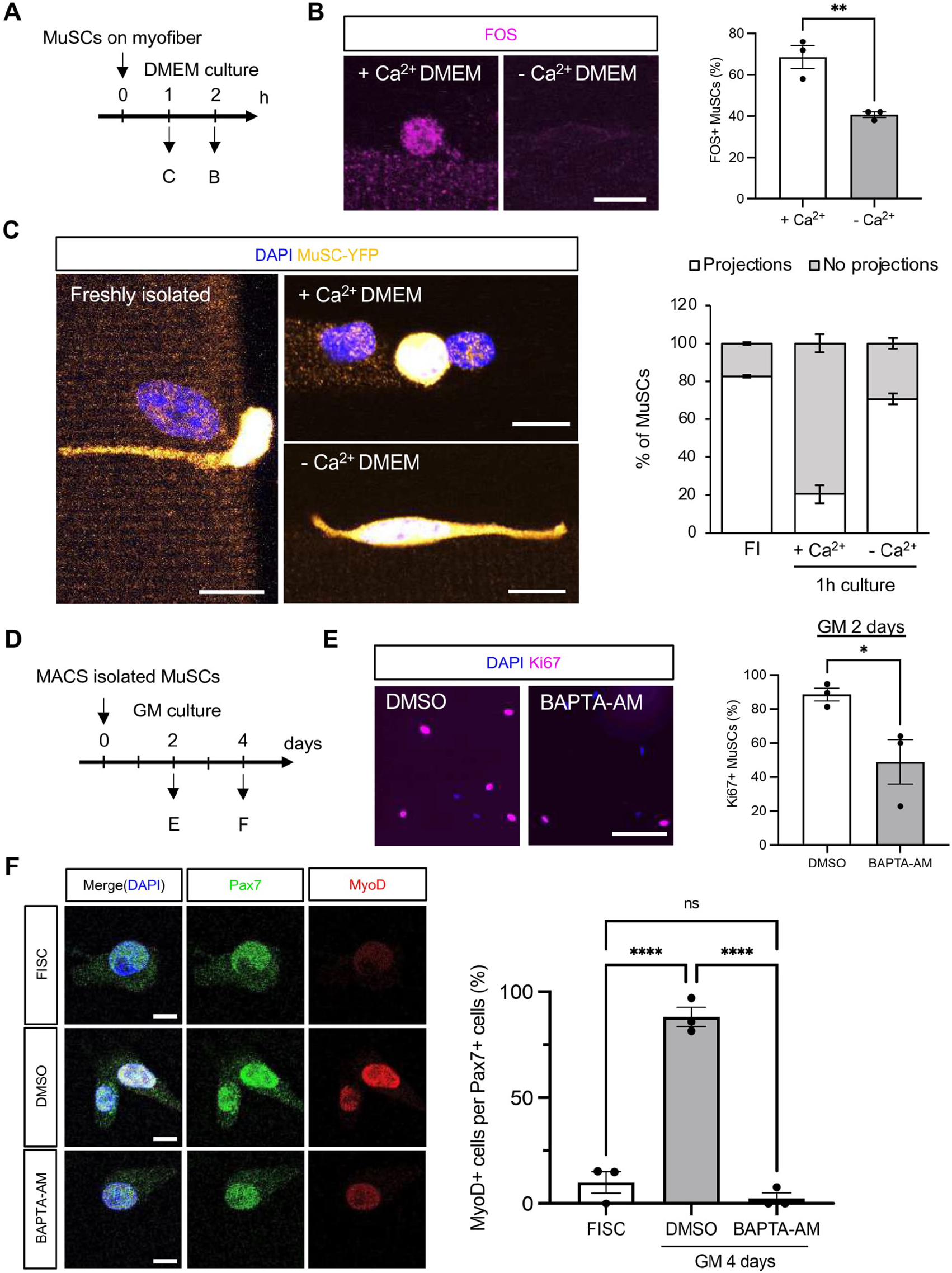
Intracellular Ca^2+^ deficiency inhibits activation of MuSC. **(A)** Time course for myofiber isolation and culture of MuSCs on myofiber. **(B)** Expression of AP-1 transcription factor of MuSCs cultured for 2 hours in DMEM with or without Ca^2+^. Left panel: Representative fluorescent images of AP-1 transcription factor (Fos; magenta). Scale bar: 10 μm. Right panel: Quantification of AP-1 expression in MuSCs ((> 50 MuSCs per condition from N = 3 mice). **(C)** Evaluation of MuSCs with quiescent projection on myofibers from Pax7CreERT2/+; Rosa26YFP. MuSCs morphology freshly isolated (FI) or cultured for 2 hours in DMEM with or without Ca^2+^. MuSC morphology was evaluated by detection of YFP (yellow). Nuclei were detected by DAPI (blue) (> 50 MuSCs per condition from N = 3 mice). Scale bar: 10 μm. **(D)** Time course for isolation and culture of MuSCs. (E)Detection of Ki67 in MuSCs 2 days after culture. The number of Ki67+ cells was counted in MuSCs in growth medium (GM) including BAPTA-AM (10 µM), intracellular Ca^2+^ chelator during 48 hours. Left: Representative images of Ki67 (megenta) and DAPI for nuclei (blue). Scale bar: 100 μm. Right: The ratio of Ki67-positive was quantified. (> 300 MuSCs per condition from N=3 mice). **(F)** Measurement of Pax7 and MyoD expression in MuSCs in culture medium with BAPTA-AM (10 µM). Left: Representative images of Pax7 (green), MyoD (red), and nuclei (blue). Scale bar: 10 μm. Percentages of MyoD-positive cells per Pax7-positive were counted in cells freshly isolated or cultured during for 4 days in GM or including 10 µM BAPTA-AM (> 40 cells per condition from N=3 mice). \**P* < 0.05, \*\**P* < 0.01, ****: *P* < 0.0001. One-way ANOVA followed by Tukey’s test.

MuSC entry into the cell cycle typically occurs within 1-2 days during *ex vivo* culture. To evaluate the involvement of Ca^2+^ signaling in this process, MuSCs were cultured for 2 days in the presence of BAPTA-AM (Fig. 4D). Immunofluorescence analysis of the proliferation marker Ki-67 revealed a significant reduction in the proportion of Ki-67-positive MuSCs under Ca^2+^ chelation (Fig. 4E), indicating impaired cell cycle entry. Finally, we examined the expression of MyoD, a key transcription factor involved in MuSC activation. MuSCs cultured for 4 days in the presence of BAPTA-AM largely remained MyoD-negative (Fig. 4F), further supporting our conclusion that Ca^2+^ signaling is necessary for the activation process. Collectively, these results demonstrate that Ca^2+^ signaling is required for multiple early steps in MuSC activation, including immediate early gene induction, morphological remodeling, and progression into the cell cycle.

### PIEZO1 mediates stiffness-dependent Ca^2+^ influx in MuSCs

In a previous study, we reported that the Ca^2+^-permeable mechanosensitive ion channel PIEZO1 is required for Ca^2+^ influx in FISCs and promotes MuSC proliferation^31^. We further demonstrated that PIEZO1 senses substrate stiffness and enhances MuSC proliferative capacity^31^. Based on these findings, we next examined whether substrate stiffness regulates PIEZO1 expression and Ca^2+^ dynamics in MuSCs. We prepared hydrogel substrates with defined stiffnesses—soft (2 kPa) and hard (32 kPa)—and cultured MuSCs on these matrices. MuSCs isolated from PIEZO1-tdTomato reporter mice were cultured for 3 days, and the fluorescence intensity of tdTomato was quantified as a readout of PIEZO1 expression (Fig. 5A). PIEZO1-tdTomato intensity did not differ significantly between MuSCs cultured on the 2 kPa and 32 kPa substrates. In contrast, the intensity was markedly increased when MuSCs were cultured on glass (∼10^6^ kPa) (Fig. 5A), suggesting that PIEZO1 expression is enhanced under extremely stiff conditions

Next, to directly assess the contribution of PIEZO1 to stiffness-dependent Ca^2+^ dynamics, we generated a MuSC-specific *Piezo1* conditional knockout (cKO) mouse model expressing the genetically encoded Ca^2+^ indicator GCaMP6s by crossing *Piezo1flox* mice with *Pax7IRES-CreERT2; Rosa26GCaMP6s* mice. *Piezo1* deletion was induced and confirmed (Supplementary Fig. 1A and 1B) by intraperitoneal tamoxifen administration for five consecutive days, followed by MuSC isolation (Fig. 5B). Isolated MuSCs were cultured on either soft (2 kPa) or hard (32 kPa) hydrogel substrates, and spontaneous Ca^2+^ dynamics were assessed by quantifying GCaMP fluorescence peaks (Fig. 5C, Supplementary Videos 1–4). In control MuSCs, culture on the stiffer substrate (32 kPa) resulted in a significant increase in GCaMP peak amplitude compared to that with cells cultured on the softer substrate (2 kPa) (Fig. 5D). Notably, this stiffness-dependent increase in Ca^2+^ signaling was abolished in *Piezo1*-deficient MuSCs, even when cultured on the 32 kPa substrates (Fig. 5D). These results indicate that PIEZO1 is required for sensing substrate stiffness and mediates stiffness-dependent spontaneous Ca^2+^ influx in MuSCs.

**Figure 5:**
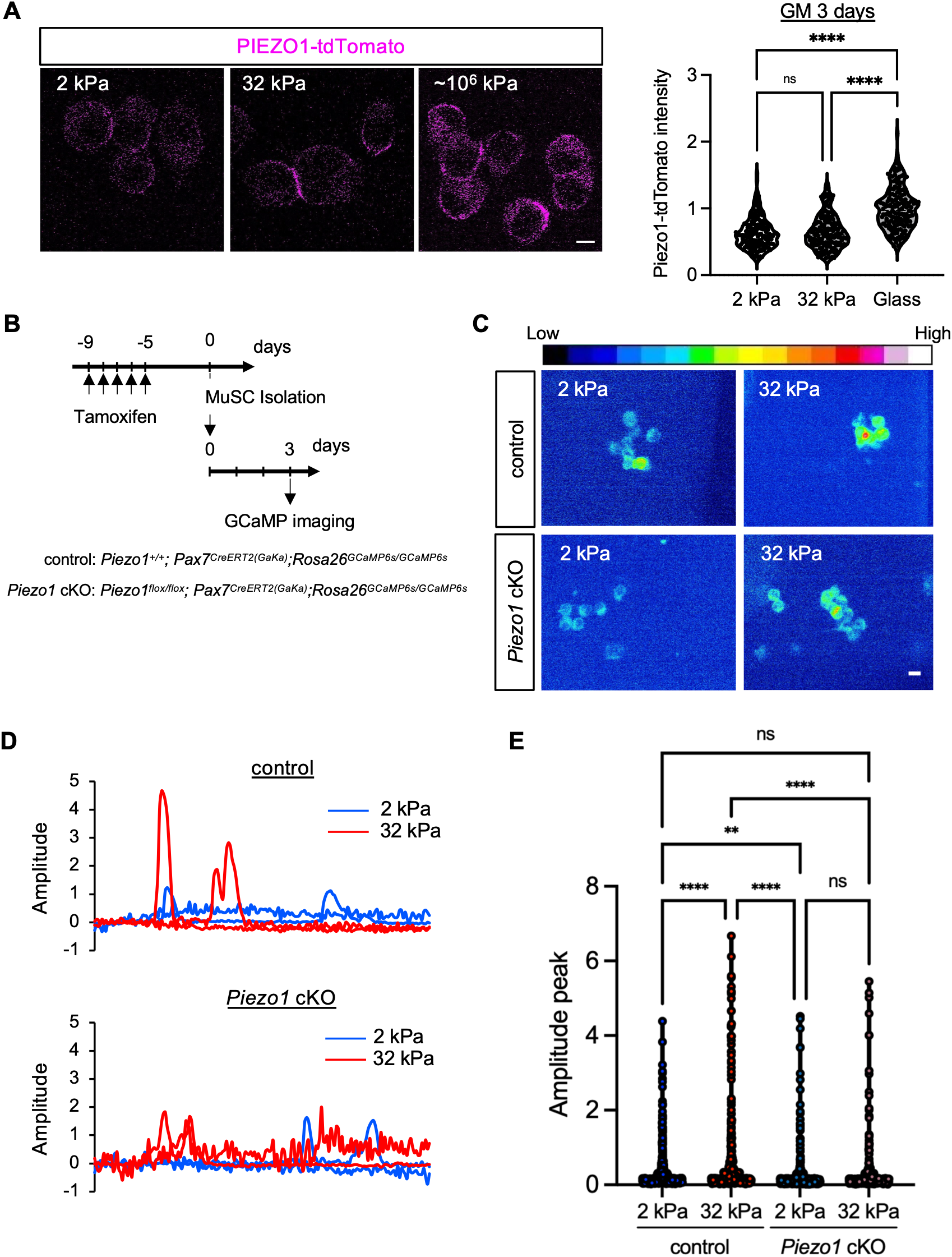
Intracellular Ca^2+^ fluctuations in MuSCs correlate with substrate stiffness. **(A)** Measurement of PIEZO1 expression level in MuSCs cultured on different elastic modulus. MuSCs were isolated from *Piezo1*^*tdTomato/tdTomato*^ mouse. Left: Detection of PIEZO1-tdTomato (magenta) in MuSCs cultured on cell culture substrate with 2 kPa, 32 kPa or glass bottom dish (∼10^6^ kPa) elastic modulus. Scale bar: 10 μm. Right: Measurement of the fluorescence intensity of PIEZO1-tdTomato (>150 cells per condition from N = 3 mice). The values were normalized relative to intensity of cells cultured on glass bottom dish. **(B)** Time course for isolation, culture and GCaMP imaging of MuSCs. **(C, D and E)** Measurement of GCaMP-based intracellular Ca^2+^ fluctuations in MuSCs cultured on different elastic modulus. **(C)** Representative fluorescence images of GCaMP6s in MuSCs cultured on cell culture substrate with 2 kPa or 32 kPa. LUTs indicate signal intensity. Scale bar: 10 μm. **(D)** Representative recording traces (2 kPa: blue trances, 32 kPa: red traces) of Amplitude from two MuSCs isolated from control and *Piezo1* cKO mice. **(E)** Quantification of the maximum amplitude during the measurement in MuSCs cultured for 3 days. GCaMP maximum peaks were defined as the maximum ΔF/F_0_ value of each calcium transient. Blue dots represent GCaMP peaks from MuSCs cultured on 2 kPa. Red dots represent GCaMP peaks from MuSCs cultured on 32 kPa. More than 100 cells from N = 3 mice per condition were investigated. \*\**P* < 0.01, \*\*\*\**P* < 0.0001. One-way ANOVA followed by Tukey’s test. See also Supplemental Videos 1 – 4.

### Mechanosensitive ion channels PIEZO1 and TRPM7 are essential for MuSC migration *in vivo* and *ex vivo*

Efficient migration of MuSCs toward sites of injury is a prerequisite for proper skeletal muscle regeneration^41-44^. Ca^2+^ signaling has been proposed as a key regulator of cell migration, driving cytoskeletal remodeling and directional movement^45^. Therefore, we investigated whether mechanosensitive ion channels mediate Ca^2+^-dependent MuSC migration. To examine the subcellular localization of PIEZO1 during MuSC migration, MuSCs isolated from PIEZO1-tdTomato reporter mice were cultured and subjected to time-lapse imaging. Notably, PIEZO1-tdTomato was enriched at the rear edge of migrating MuSCs, corresponding to the trailing end of the cell during directional movement (Fig. 6A, Supplementary Video 5).

**Figure 6:**
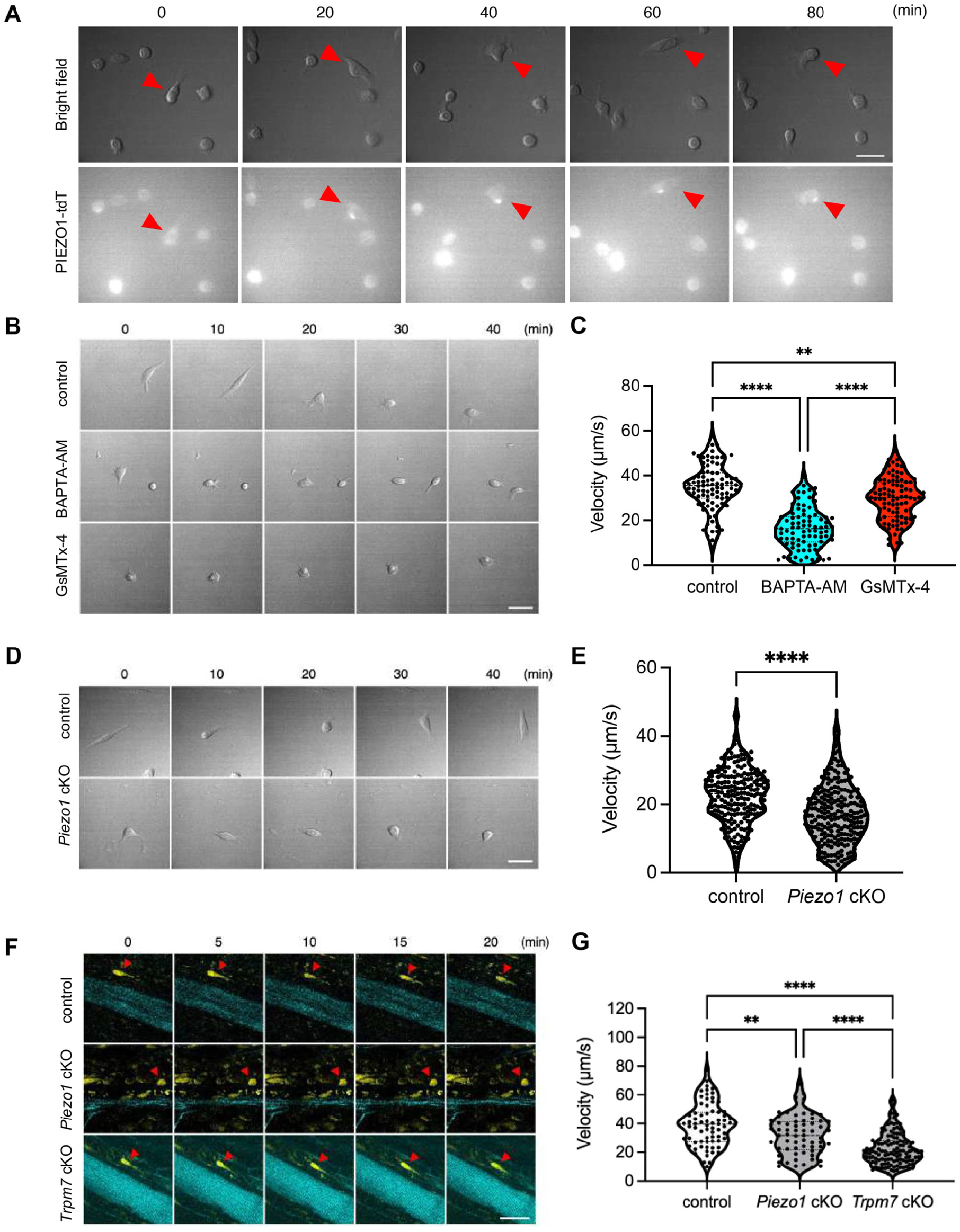
Mechanosensitive Ca^2^+ signalling controls MuSC migration during skeletal muscle regeneration. **(A)** Time-lapse imaging of MuSCs isolated from *Piezo1*^*tdTomato/tdTomato*^ mouse. Red arrow indicates the localization of PIEZO1-tdTomato at the rear end of MuSC during cell migration. Upper panel: Bright field. Lower panel: PIEZO1-tdTomato signals. Scale bar: 20 μm. See also Supplemental Video 5. **(B and C)** *Ex vivo* analysis of MuSC migration. **(B)** Representative migration trajectories of MuSCs treated with BAPTA-AM (20 µM) or GsMTx4 (300 nM) to assess the contribution of intracellular Ca^2^+ signaling and mechanosensitive ion channels to MuSC migration. Scale bar: 20 μm. **(C)** Quantification of migration speed of MuSCs under control conditions and following treatment with BAPTA-AM (blue) or GsMTx4 (red). See also Supplemental Video 6 – 8. **(D and E)** *Ex vivo* analysis of *Piezo1*-deficient MuSC migration. **(D)** Representative migration trajectories of MuSCs isolated from control and *Piezo1*-deficient mice under *ex vivo* conditions. Scale bar: 20 μm. **(E)** Quantification of MuSC migration speed in control and *Piezo1*-deficient MuSCs. White violins indicate control MuSCs, and gray violins indicate *Piezo1* cKO MuSCs. See also Supplemental Video 9 and 10. **(F and G)** *In vivo* analysis of MuSC migration during muscle regeneration. **(F)** Representative images obtained from control, *Piezo1*-deficient, and *Trpm7*-deficient mice. Two-photon excitation enabled visualization of YFP-labeled MuSCs, while collagen fibrils were simultaneously imaged via second harmonic generation (SHG) signals. Scale bar: 50 μm. **(G)** Quantification of *in vivo* MuSC migration speed. White violins indicate control mice, gray violins indicate *Piezo1* cKO mice, and dark gray violins indicate *Trpm7* cKO mice. \*\**P* < 0.01, \*\*\*\**P* < 0.0001. One-way ANOVA followed by Tukey’s test. See also Supplemental Video 11 – 13.

Next, we quantified MuSC migration velocity under Ca^2+^ perturbation. Chelation of intracellular Ca^2+^ using BAPTA-AM significantly reduced MuSC migration speed, highlighting the importance of Ca^2+^ signaling in MuSC motility (Fig. 6B and 6C, Supplementary Videos 6 and 7). Similarly, treatment with GsMTx-4, a peptide inhibitor of mechanosensitive ion channels including PIEZO1, resulted in reduced migration velocity, indicating that Ca^2+^ influx mediated by mechanosensitive ion channels contributes to MuSC migration (Fig. 6B and 6C, Supplementary Video 8).

Based on these results, we hypothesized that PIEZO1 directly regulates MuSC migration. To test this, MuSCs isolated from MuSC-specific *Piezo1*-cKO mice were cultured *ex vivo*, and their migration velocity was quantified. *Piezo1*-deficient MuSCs exhibited significantly reduced migration speed compared to that in control cells (Fig. 6D and 6E, Supplementary Videos 9 and 10), indicating that PIEZO1 is required for efficient MuSC migration *ex vivo*.

To further validate these findings, we performed intravital two-photon microscopy to monitor MuSC migration in injured muscle (Supplementary Fig. 2A and 2B). MuSC from *Piezo1*-cKO mice that underwent muscle injury for 3 days exhibited markedly reduced migration velocity *in vivo*, consistent with the *ex vivo* observations (Fig. 6F and 6G, Supplementary Videos 11 and 12).

Finally, we examined the involvement of another mechano-sensitive ion channel, TRPM7, which we recently identified as a regulator of MuSC activation and muscle regeneration^39^. Following confirmation of *Trpm7* deletion (Supplementary Fig. 2C and 2D), intravital imaging of MuSCs from *Trpm7* cKO mice 3 days following muscle injury showed a similar reduction in their migration velocity (Fig. 6F and 6G, Supplementary Videos 11 and 13), indicating that TRPM7 also contributes to MuSC migration during muscle regeneration. Collectively, these results demonstrate that the mechano-sensitive ion channels PIEZO1 and TRPM7 are essential regulators of Ca^2+^-dependent MuSC migration both *ex vivo* and *in vivo*.

## Discussion

The pioneering discoveries of Dr. Setsuro Ebashi established Ca^2+^ as a central regulator of skeletal muscle, laying the foundation for the field of excitation–contraction coupling ^46-49^. In contrast, the contribution of Ca^2+^ signaling in MuSC has remained relatively understudied. Previous studies, including our own, have suggested a role for Ca^2+^ signaling in MuSC proliferation^31^ and myogenic differentiation^25^; however, the regulation of Ca^2+^ dynamics during early MuSC activation has not been fully elucidated.

In this study, we demonstrate that spontaneous Ca^2+^ influx is a prominent feature of activated and proliferative MuSCs across multiple physiological and pathological contexts, including muscle regeneration, dystrophy, aging, and *ex vivo* activation. These findings position Ca^2+^ dynamics as a core biophysical signature associated with MuSC activation states. Compared with our previous studies^31,39^, in which MuSC Ca^2+^ dynamics were mainly examined on glass substrates, the present study analyzed Ca^2+^ dynamics on substrates with stiffness closer to that of skeletal muscle during regeneration. This approach allowed us to assess MuSC Ca^2+^ dynamics under a mechanical environment that more closely resembles *in vivo* conditions. In addition, the MACS-based isolation method enabled rapid and high-yield isolation of MuSCs, allowing live-cell Ca^2+^ imaging to be performed under conditions closer to the freshly isolated state. A major technical challenge in studying ion dynamics in MuSCs has been the difficulty of isolating cells while preserving their physiological state. By combining optimized MuSC isolation strategies with time-resolved flow cytometry and live-cell Ca^2+^ imaging using both chemical Ca^2+^ indicators and a genetically encoded Ca^2+^ sensor, our workflow enables quantitative assessment of Ca^2+^ dynamics at single-cell resolution (Fig. 1 and 2). Notably, this approach led us to investigate Ca^2+^ dynamics across diverse MuSC contexts, including MuSCs derived from fast- and slow-twitch muscle fibers, dystrophic muscles, aged skeletal muscles, and regenerating tissues, thereby enabling a comparative analysis of Ca^2+^ signaling under distinct physiological and pathological conditions (Fig. 3). While further validation is required, Ca^2+^ signaling profiles could potentially be exploited as biophysical biomarkers to distinguish quiescent, activated, and dysfunctional MuSC populations across regeneration, aging, and disease contexts. Importantly, this platform is not limited to Ca^2+^ signaling. Advances in fluorescent probes^50,51^ and genetically encoded sensors^52-54^ have enabled the measurement of additional intracellular biophysical parameters. Such probes, including intracellular pH sensors and thermosensitive indicators developed in previous studies, could be readily integrated into our workflow. This could allow systematic investigation of how multiple biophysical cues, such as ion flux, pH regulation, and intracellular thermal changes, collectively influence MuSC behavior during regeneration.

One notable observation in our study is that spontaneous Ca^2+^ fluctuations were readily detectable in *ex vivo*-activated MuSCs but were less apparent in MuSCs directly isolated following CTX-induced injury (Fig. 1 and 2). This discrepancy likely reflects differences in the temporal resolution and mechanical environment of MuSCs *in vivo* compared to that *in vitro. In vivo*, activated MuSCs reside within a highly constrained niche, are tightly associated with the extracellular matrix (ECM), and are exposed to various physiological molecules secreted by surrounding cells. Under these conditions, Ca^2+^ signaling events may be rapid, spatially restricted, or asynchronous across the population, making them difficult to capture using bulk isolation-based approaches. In contrast, *ex vivo* culture provides a permissive mechanical and biochemical environment that amplifies spontaneous Ca^2+^ dynamics, enabling their detection and quantitative analysis. These findings highlight the importance of experimental context when interpreting ion signaling events in stem cells.

In this study, we demonstrate that spontaneous Ca^2+^ influx peaks approximately 2 days after MuSC activation (Fig. 1 and 2), coinciding with robust cell-cycle entry and mitotic activity^55^. Notably, PIEZO1 has previously been reported to localize to the midbody during cytokinesis, suggesting a potential role for Ca^2+^ regulation during cell division^31,56^. Consistent with this, we observed enhanced Ca^2+^ dynamics during the stage of active MuSC proliferation. Importantly, pharmacological inhibition of Ca^2+^ signaling using the intracellular Ca^2+^ chelator BAPTA-AM markedly attenuated multiple hallmarks of MuSC activation, including induction of immediate early genes, morphological remodeling, cell-cycle entry, and MyoD expression (Fig. 4). These findings provide functional evidence that Ca^2+^ signaling is not merely correlated with MuSC activation but is essential for progression through early activation and proliferative phases. Collectively, these results raise the possibility that Ca^2+^ influx contributes not only to the initiation of MuSC activation but also to the coordination of cell-cycle progression and cytokinetic events. Whether Ca^2+^ signals directly regulate mitotic machinery or act indirectly through cytoskeletal remodeling and mechanotransduction pathways remains an important question for future investigation.

In this study, we also demonstrate that Ca^2+^ fluctuations are more pronounced under certain conditions, including in dystrophic mdx mice and aged mice (Fig. 3). As muscle degeneration and insufficient regeneration are hallmarks of mdx mice, MuSCs residing in dystrophic muscle are prone to aberrant activation, thereby recapitulating features observed in MuSCs isolated from CTX-injured muscle. In addition to the enhanced degeneration–regeneration cycles observed in mdx mice, the microenvironmental niche surrounding MuSCs is markedly altered in both dystrophic and aged muscles. Notably, increased tissue stiffness resulting from changes in ECM composition, such as elevated collagen content, has been reported in aged animals and dystrophic models. For example, direct measurements using atomic force microscopy have demonstrated that aged muscle is stiffer than young muscle^57^. Although aged MuSCs showed elevated Ca^2+^ fluctuations (Fig. 3I–3L), this increase should not necessarily be interpreted as enhanced regenerative capacity. Aging-associated changes in tissue mechanics have been shown to influence adult stem/progenitor cell function; for example, stiffening of the aged central nervous system niche impairs oligodendrocyte progenitor cell function through mechanosensitive signaling involving PIEZO1^58^. In our study, PIEZO1-tdTomato fluorescence intensity was not significantly altered in MuSCs isolated from aged PIEZO1-tdTomato mice at 3 days after CTX-induced injury compared with young controls (Supplementary Fig. 3).Thus, the elevated Ca^2+^ fluctuations in aged MuSCs may not be attributable simply to increased PIEZO1 expression, but may instead reflect aging-associated changes in mechanotransduction, membrane mechanics, cytoskeletal organization, or Ca^2+^ handling.

The mechanisms by which MuSCs sense altered biophysical properties associated with muscular dystrophy and aging remain unclear. One plausible explanation is the involvement of mechanosensory and/or mechanotransduction systems. Consistent with this hypothesis, our results show increased PIEZO1-tdTomato expression and enhanced Ca^2+^ fluctuations under high-stiffness conditions. Importantly, this stiffness-dependent increase in Ca^2+^ fluctuations was abolished in *Piezo1*-deficient MuSCs (Fig. 5D), suggesting that PIEZO1 is a key component that mediates Ca^2+^ fluctuations in MuSCs by sensing changes in the mechanical environment. Further studies, including identification of Ca^2+^-dependent downstream signaling pathways, will be required to elucidate the functional significance of Ca^2+^ fluctuations in MuSCs.

Efficient migration of MuSCs toward sites of injury is essential for effective muscle regeneration^42,44^. The mechanical properties of the MuSC niche, including ECM stiffness, dynamically change during regeneration and are known to influence MuSC behavior. In this study, we demonstrate that PIEZO1 functions as a stiffness-sensitive regulator of Ca^2+^ dynamics in MuSCs. Although PIEZO1 expression was comparable on soft (2 kPa) and intermediate (32 kPa) hydrogel substrates, it was markedly upregulated on extremely stiff substrates such as glass (Fig. 5A). Importantly, despite this minimal change in expression between 2 and 32 kPa, MuSCs could sense these subtle differences in substrate stiffness and exhibited enhanced Ca^2+^ dynamics on the stiffer substrate in a PIEZO1-dependent manner (Fig. 5C and 5D). These findings indicate that PIEZO1 primarily functions as a mechanosensitive Ca^2+^ gate that translates fine-scale mechanical differences within a physiologically relevant stiffness range into intracellular Ca^2+^ signaling, rather than solely through modulation of its expression levels.

We previously reported that mechanosensitive ion channels, including PIEZO1 and TRPM7, regulate cytoskeletal rearrangements during MuSC activation^31,39^. In this study, we extend these findings by demonstrating that both channels are required for efficient MuSC migration *in vivo* and *ex vivo* (Fig. 6). Notably, PIEZO1 was enriched at the rear edge of migrating MuSCs in this study, whereas TRPM7 has been reported to localize toward the leading edge in other migrating cell types^59^. This spatial segregation suggests that distinct mechanosensitive ion channels may play complementary roles during cell migration, coordinating rear-end retraction and front-end protrusion through localized Ca^2+^ signaling and mechanotransduction^45^. Because directed migration in response to substrate stiffness, known as durotaxis^60^, requires spatially coordinated mechanosensing, PIEZO1- and TRPM7-associated signaling may contribute to the polarized responses of MuSCs during migration. Notably, emerging evidence indicates that intracellular signaling in adult stem cells is not governed by Ca^2+^ alone but rather by the coordinated actions of multiple cations, including Mg^2+^ and Zn^2+^, thereby contributing to metabolic regulation, enzyme activity, and transcriptional control^61^. This division of labor among ion channels may enable MuSCs to translate mechanical cues into directed movement during regeneration.

In summary, our findings establish Ca^2+^ dynamics as a fundamental feature of MuSC activation, proliferation, and migration. By integrating genetic, pharmacological, and biophysical approaches, the mechanosensitive ion channel PIEZO1 act as key transducers linking mechanical environments to intracellular Ca^2+^ signaling. This study provides a conceptual framework for understanding how biophysical signals regulate adult stem cell behavior and opens new avenues for targeting ion-based mechanisms in muscle regeneration and disease.

## Acknowledgements

We thank members of the Hara laboratory for their scientific contributions. This study was supported by the Grant-in-Aid for Scientific Research KAKENHI (22H03484, 25K02994); Grant-in-Aid for Transformative Research Areas B (23H03854; 23H03856); Intramural Research Grant (5-6, 8-13) for Neurological and Psychiatric Disorder of NCNP; grants from Takeda Science Foundation, Uehara memorial foundation, Chugai Foundation for Innovative Drug Discovery Science (SRG2022), the Asahi Glass Foundation and an internal grant from the University of Shizuoka to Y Hara; and Grant-in-Aid for JSPS Research Fellow (20J22984), Grant-in-Aid for Scientific Research KAKENHI (23K19898, 24K20602) and an internal grant from the University of Shizuoka. to K.Hirano; and Grant-in-Aid for JSPS Research Fellow (23KJ1814) to Y.Ishikawa. This work was supported by Kyoto University Live Imaging Center. This work was supported in part by Grants-in-Aid KAKENHI 16H06280 “ABiS”.

## Author contributions

Conceptualization: K.H., Y.I., and Y.H.; Methodology: N.M, K.Y., A.M., and M.T., Visualization: K.H., and Y.H.; Investigation: K.H., Y.I., Y.K., H.W., M.S., T.Y., Y.Y., Yu.A., Supervision: Y.O, K.N., Yo.A., Writing—original draft: K.H., and Y.H.; Writing—review & editing: K.H., Y.I., and Y.H.

## Materials and Methods

### Mice

Animal care, ethical use, and protocols were approved by the Animal Care Use and Review Committee of the University of Shizuoka (#256739, #256740, #256742). The protocol includes justification of the use of animals, their welfare, and the incorporation of the principles of the “3Rs” (Replacement, Reduction, and Refinement). Wild-type mice were maintained on a C57BL/6J background. The mdx mice used in this study were maintained as an inbred mdx line after crossing the original B10 mdx mice with C57BL/6J mice for at least five to six generations. *Piezo1*^*flox*^ mice were purchased from the UC Davis KOMP repository as previously described^31^. *Pi-ezo1*^*flox*^ mice were further mated with *Pax7*^*CreERT2/+(Fan)*^ transgenic mice^62^ (The Jax laboratory, strain ID: 012476) or *Pax7*^*IRESCreERT2/+(Kardon)*^ transgenic mice^33^ (The Jax laboratory, strain ID: 017763) to generate MuSC-specific *Piezo1*-deficient mice. *Piezo1* cKO (*Pax7*^*IRESCreERT2/+(Kardon)*^) was used in Fig. 5 and *Piezo1* cKO (*Pax7*^*CreERT2/+(Fan)*^) was used in Fig. 6. *Rosa26GCaMP6s* mice were obtained from RIKEN. The primers used for genotyping are listed in Supplementary Table 2. Tamoxifen (Sigma; dissolved in corn oil at a concentration of 20 mg/mL) was used to induce Cre recombinase expression. The mice were injected intraperitoneally with tamoxifen with 2 mg tamoxifen daily for 5 consecutive days. 50 µL and 100 µL of 10 µM CTX (#L8102, Latoxan) were respectively injected into the tibialis anterior and gastrocnemius muscle of mice.

### MuSCs isolation using MACS

MuSCs from the limb muscles were isolated as previously described^16^. Skeletal muscle samples obtained from the hindlimbs of mice were subjected to collagenase treatment using 0.2% collagenase type II (Worthington Biochemical). Mononuclear cells were incubated with phycoerythrin (PE)-conjugated anti-mouse Ly-6A/E (Sca-1) antibody (#122508, BioLegend), PE-conjugated anti-mouse CD45 antibody (#110708, BioLegend), PE-conjugated anti-mouse CD31 antibody (#102508, BioLegend) and anti-mouse Integrin-α7 antibody (#K0046-3, MBL) at 4°C for 30 min, followed by Anti-PE magnetic MicroBeads (#130-048-801, Miltenyibiotec). Sca-1-/CD45-/CD31-cells were collected by magnetic bead separation on the autoMACS® Pro Separator (Miltenyibiotec). These cells were treated with Anti-Mouse IgG magnetic MicroBeads (#130-048-402, Miltenyibiotec). Target cells (Integrin-α7+ cells) were collected by magnetic bead separation on the autoMACS®.

### Calcium imaging (Cal-520)

Cytosolic Ca^2+^ was detected by fluorogenic calcium-sensitive dye Cal-520 (#21130, AAT Bioquest). MACS isolated MuSCs were incubated with Brilliant Violet 421 (BV421)-conjugated anti-Mouse CD106 (VCAM-1) antibody (#740019, BD Bioscience) at 4°C for 1 hr. These cells were sorted into 1.5-ml tubes containing growth medium (GM) (DMEM supplemented with 30% fetal bovine serum, 1% chicken embryo extract (US Biological), 10 ng/mL basic fibroblast growth factor (ORIENTAL YEAST Co., Ltd.), and 1% penicillin-streptomycin). Then, 10 µM calcium indicator Cal-520, AM, and 0.02% Pluronic F-127 (#P2443-250G, Sigma-Aldrich) were added into the sorted tubes before incubating for 30 min at 37°C (5% CO2). The stained MuSCs were sorted with SONY MA900 and detected with fluorescein isothiocyanate (FITC) for analysis.

### Calcium imaging (Fura-2)

Fura-2 imaging was performed as previously described with minor modifications^31^. Cytosolic Ca^2+^ was detected by fluorogenic calcium-sensitive dye Fura-2 AM (#343-05401, Dojindo). For indicator loading, MuSCs were plated on glass-bottomed dishes (#D11531H, Matsunami) coated with Matrigel and incubated with 5 µM Fura-2 AM in growth medium at 37°C for 30 min (5% CO2). Time-lapse images were obtained every 2 s using an epifluorescence microscope (Axio Observer.Z1; Zeiss) equipped with a 20× dry objective lens (NA 0.5). The base composition of HBS was (in mM) 140 NaCl, 5 KCl, 2 MgCl_2_, 2 CaCl_2_, 10 glucose, and 10 HEPES (pH = 7.4 adjusted with NaOH). Ratiometric images (F340/F380) were analyzed using Physiology software (Zeiss). These experiments were performed using a heat chamber (Zeiss) to maintain the temperature at 37°C throughout the imaging process. The amplitude was calculated using the following formula:

Amplitude = [value of the F340/F380 ratio] - [average value of the F340/F380 ratio during the measurement]

The Peak amplitude was defined as the maximum amplitude value observed during the recording for each cell, and this value was used for the quantification.

### Calcium imaging (GCaMP)

Cytosolic Ca^2+^ was detected by the genetically encoded calcium indicator GCaMP6s. MuSCs were isolated from Rosa26GCaMP6s mice and plated on glass-bottomed dishes coated with Matrigel and incubated with HBS at 37°C for 15 min (5% CO2). Time-lapse images were obtained every 2 s using an epifluorescence microscope (Axio Observer.Z1; Zeiss) equipped with a 20× dry objective lens (NA 0.5). Ratiometric images (F/F0) were analyzed using Physiology software (Zeiss). These experiments were performed using a heat chamber (Zeiss) to maintain the temperature at 37°C throughout the imaging process. The amplitude was calculated using the following formula:

Amplitude = [value of the GCaMP intensity] - [average value of the GCaMP intensity during the measurement]

The Peak amplitude was defined as the maximum amplitude value observed during the recording for each cell, and this value was used for the quantification.

### Single myofiber isolation

Myofibers were isolated from the Extensor Digitorum Longus (EDL) muscles, as previously described^31^. Isolated EDL muscle samples were incubated with 0.2% collagenase I (Sigma-Aldrich) in DMEM at 37 °C for 2 h. Myofibers were released by gently flushing the muscle samples in plating medium using a fire-polished glass pipette. For Fig. 4A-C, myofibers were prepared using a modified protocol^20^. Isolated EDL muscle samples were incubated with 0.2% collagenase I (#C0130, Sigma-Aldrich) prepared in Ca^2+^-free DMEM (Gibco, Thermo Fisher Scientific) containing 20 µM BAPTA-AM (B035, Dojindo) at 37°C for 1 h. Myofibers were then released by gently flushing the muscle samples using a fire-polished glass pipette. The isolated myofibers were cultured in Ca^2+^-free DMEM supplemented with 20 µM BAPTA-AM.

### MuSC culture

MuSCs were cultured in growth medium (DMEM supplemented with 30% foetal bovine serum (Sigma-Aldrich), 1% chicken embryo extract (US Biological), 10 ng/ml basic fibroblast growth factor (ORIENTAL YEAST Co., Ltd.), and 1% penicillin–streptomycin (FUJI-FILM Wako Pure Chemical Corporation)) on culture dishes coated with Matrigel (Corning). For hard substrates, glass-bottom dishes (#D11531H, Matsunami) were used. For stiffness-tuned substrates, commercially available CytoSoft® plates with polydimethylsiloxane (PDMS) layers were used. The elastic moduli of the substrates were 2 kPa (#5185-1EA, Advanced BioMatrix) and 32 kPa (#5188-1EA, Advanced BioMatrix). CytoSoft® substrates were coated with PureCol® collagen solution (#5005-100ML, Advanced BioMatrix) prior to cell seeding according to the manufacturer’s instructions.

### Immunofluorescent analysis

The cells were placed in Matrigel-coated glass-bottom dishes and fixed with 4% paraformaldehyde (PFA)/PBS for 10 min. After permeabilization in 0.1% Triton X-100/PBS for 10 min, the samples were blocked in 1% BSA/PBS for 1 h and probed with the antibodies listed in Supplementary Table 1 at 4°C overnight. After multiple washes with PBS, the secondary antibodies listed in the Supplementary Table 1 were added. Nuclei were detected using DAPI (Dojindo, 1:1000). Myofibers were fixed with 2% PFA/PBS for 5 min. After permeabilization and blocking with 0.1% Triton X-100 in 1%BSA/PBS for 15 min, the samples were probed with the antibodies listed in the Supplementary Table 2 at room temperature for 2 h or at 4°C overnight. After washing once with PBS, the secondary antibodies listed in the Supplementary Table 2 were added. Nuclei were detected using DAPI (Dojindo, 1:1000). Immunofluorescent signals were visualized with Alexa488- or Alexa555-conjugated secondary antibodies using an epifluorescence microscope (Axio Observer.Z1, Zeiss) equipped with 10× dry, 20× dry, and 40× dry objective lenses with numerical apertures of 0.5, 0.5, and 0.75, respectively or a confocal microscope (LSM 800, Zeiss) with a 63×oil-immersion objective lens (NA 1.4). The fluorescence intensity was quantified using ImageJ software for statistical analyses.

### Histological analysis

Cardiotoxin experiments were performed as previously described. Fifty microliters of 10 µM cardiotoxin (latoxan) were injected into the tibialis anterior muscle of 8-to 15-week-old mice. The muscle was harvested at the time points indicated in each figure and snap-frozen in isopentane cooled with liquid nitrogen. Cross-cryosections (thickness, 7 µm) of the muscle samples were used for haematoxylin and eosin staining, as previously described.

### Time-lapse imaging of *ex vivo* MuSC migration

MuSCs were cultured in growth medium, which was refreshed immediately prior to imaging. Time-lapse imaging was performed using a fluorescence microscope (LCV110; Olympus). Images were acquired at 40× magnification under the following conditions: exposure time of 30 ms for DIC images and 1,000 ms for RFP fluorescence when imaging PIEZO1-tdTomato signals. Images were captured at 10-min intervals for a total duration of 72 h. Acquired image stacks were saved as stack files (. stk format) for subsequent analysis.

### Intravital imaging of MuSC migration

*Pax7*^*CreERT2*^; *Rosa26*^*YFP*^ mice were used for intravital imaging of MuSCs. To induce YFP labeling in MuSCs, tamoxifen was administered by intraperitoneal injection at a concentration of 2 µg/mL (100 µL per injection) once daily for five consecutive days. Five days after the final tamoxifen injection, muscle injury was induced by intramuscular injection of cardiotoxin (10 µM, 50 µL) into both tibialis anterior muscles. Three days after injury, *in vivo* imaging was performed using a two-photon fluorescence microscope (FV1000MPE-IX83; Olympus) as previously described^41^. During imaging, mice were maintained under anesthesia by inhalation of isoflurane. Images were acquired using a 30× silicone oil-immersion objective lens (UPLSAPO 30XS, NA 1.05, working distance 0.8 mm; Olympus) with an excitation wavelength of 940 nm. Z-stack images were collected with a step size of 0.5 µm, covering 21–33 optical sections along the z-axis. Time-lapse imaging was performed at 5-min intervals for a total duration of 4–5 h.

### Migration analysis and cell tracking

Quantitative analysis of MuSC migration was performed using ImageJ, based on previously reported methods^63^. Stack image data were imported into ImageJ and processed as follows. Image brightness and contrast were adjusted uniformly across samples. Images were cropped to a 375 × 375-pixel square region to exclude non-informative dark areas. Noise reduction was applied using a median filter with a radius of 3.0 pixels. Cell tracking was performed using the TrackMate plugin. Calibration settings were set with pixel width, height, depth, and interval all defined as 1.0. Cropping parameters were defined as X: 0–374, Y: 0–374, Z: 0–0, and T: 0–144. Spot detection was conducted using the LoG detector with an estimated blob diameter of 20 pixels and a threshold of 2.0. Median filtering and sub-pixel localization were disabled. Detected spots were visualized using the Hyper-Stack Displayer. Spot quality thresholds were manually adjusted to label individual cells. Tracking was performed using the Simple LAP tracker, ignoring cell division and fusion events, with the following parameters: linking maximum distance of 60 pixels, gap-closing maximum distance of 60 pixels, and a maximum frame gap of 2. Tracks containing fewer than seven spots were excluded from further analysis. Track and spot statistics were exported and saved as Excel files (“Migration_Data.xlsx”), with validated track statistics and spot statistics stored in separate worksheets for down-stream quantitative analysis.

### Statistical analysis

Statistical analyses were performed using Microsoft Excel or JMP 11 (JMP Statistical Discovery LLC). The statistical significance of the differences between the mean values was analyzed using a non-paired *t*-test (two-sided). One-way ANOVA followed by Tukey’s multiple-comparisons test was performed. P values of \**P* < 0.05, \*\**P* < 0.01, \*\*\**P* < 0.001 **** and *P* < 0.0001 were considered statistically significant. The results are presented as the mean + SEM. n.s. indicates results that are not statistically significant.

---

**Figure S1:**
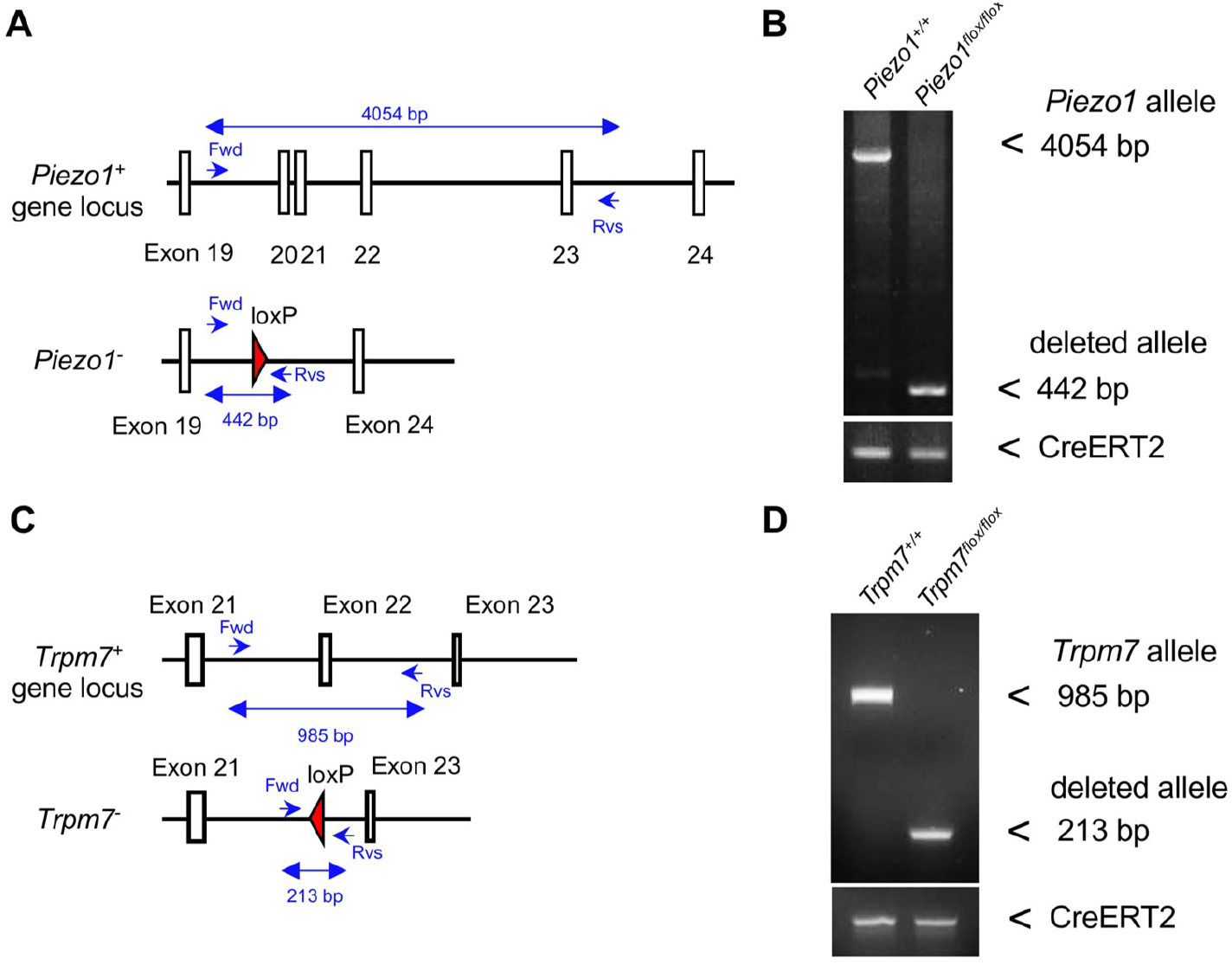
Conditional knockout of *Piezo1* and *Trpm7* in MuSC. **(A)** Map showing PCR primers for detecting deleted *Piezo1* alleles (Exon 20 – Exon 22) from wild type *Piezo1* alleles and deleted *Piezo1* alleles. **(B)** Genomic PCR detection of *Piezo1* and deleted *Piezo1* alleles in MuSCs isolated from *Piezo1*^*+/+*^; *Pax7*^*CreERT2/+*^ (con) and *Piezo1*^*flox/flox*^; *Pax7*^*CreERT2/+*^ (cKO) using the forward (Fwd) and reverse (Rvs) primers from (A). The presence of the *Pax7*^*CreERT2*^ allele in both control and *Piezo1* cKO samples was confirmed by a separate PCR detecting the CreERT2 transgene (lower panel). **(C)** Map showing PCR primers for detecting deleted *Trpm7* alleles (Exon 22) from wild type *Trpm7* alleles and deleted *Trpm7* alleles. **(D)** Genomic PCR detection of *Trpm7* and deleted *Trpm7* alleles in MuSCs isolated from *Trpm7*^*+/+*^ *Pax7*^*CreERT2/+*^ (con) and *Trpm7*^*flox/flox*^ *Pax7*^*CreERT2/+*^ (cKO) using the forward (Fwd) and reverse (Rvs) primers from (C). The presence of the *Pax7*^*CreERT2*^ allele in both control and *Trpm7* cKO samples was confirmed by a separate PCR detecting the CreERT2 transgene (lower panel).

**Figure S2:**
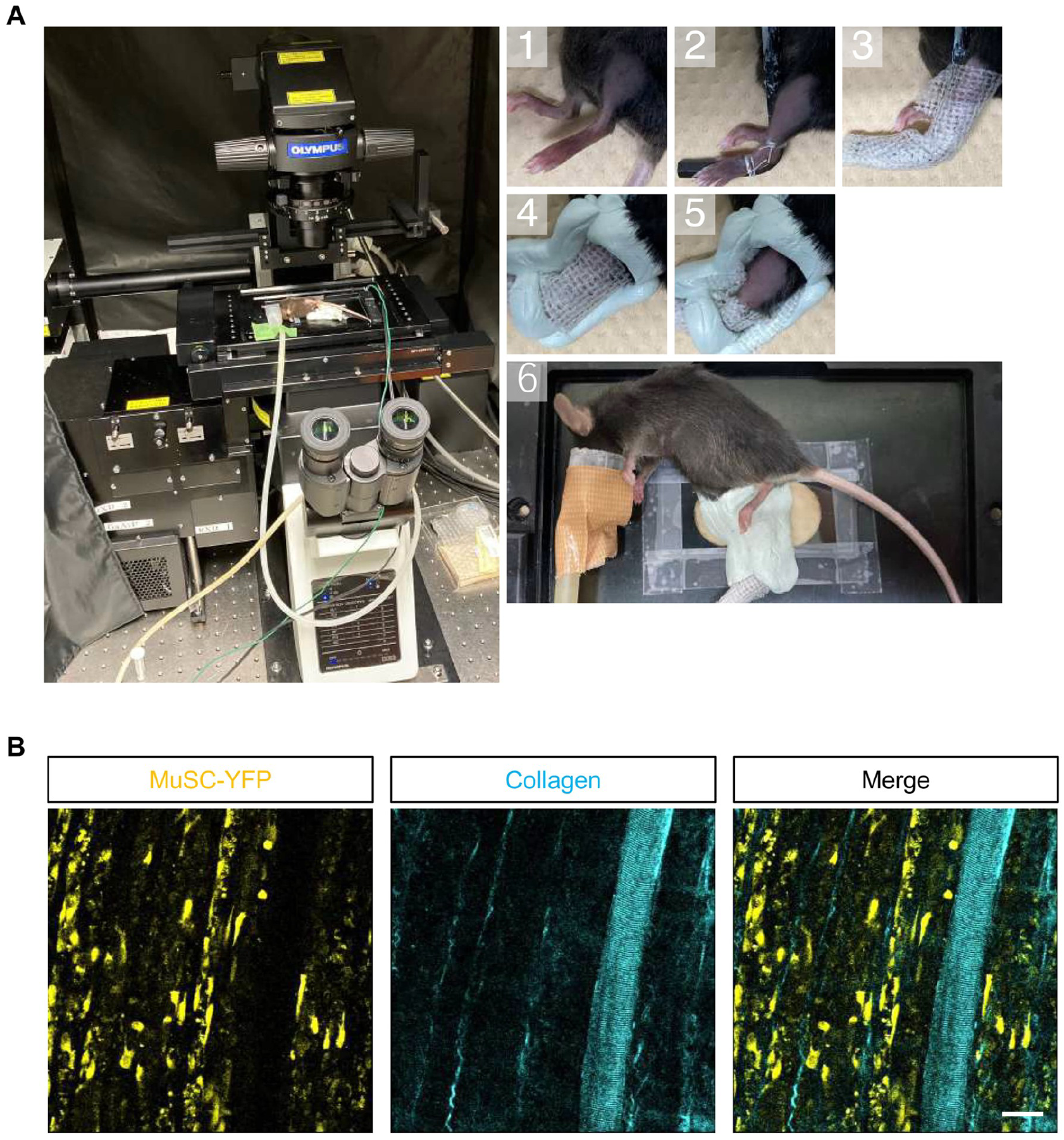
Intravital imaging of MuSCs during muscle regeneration. **(A)** Schematic overview of the intravital two-photon fluorescence microscopy setup and mouse preparation procedure. Steps 1–3 illustrate fixation of the mouse hindlimb using a hex wrench. In step 4, a simple cast is formed using epoxy putty to stabilize the limb. In steps 5 and 6, the skin over the region of interest is carefully removed to expose the underlying muscle. In step 7, the exposed muscle is immobilized on a glass plate for imaging. **(B)** Representative intravital time-lapse imaging of MuSC dynamics at 3 days after muscle injury. MuSCs are labelled with YFP. Collagen fibrils are visualized by second harmonic generation (SHG) signals. Scale bar: 50 μm.

**Figure S3:**
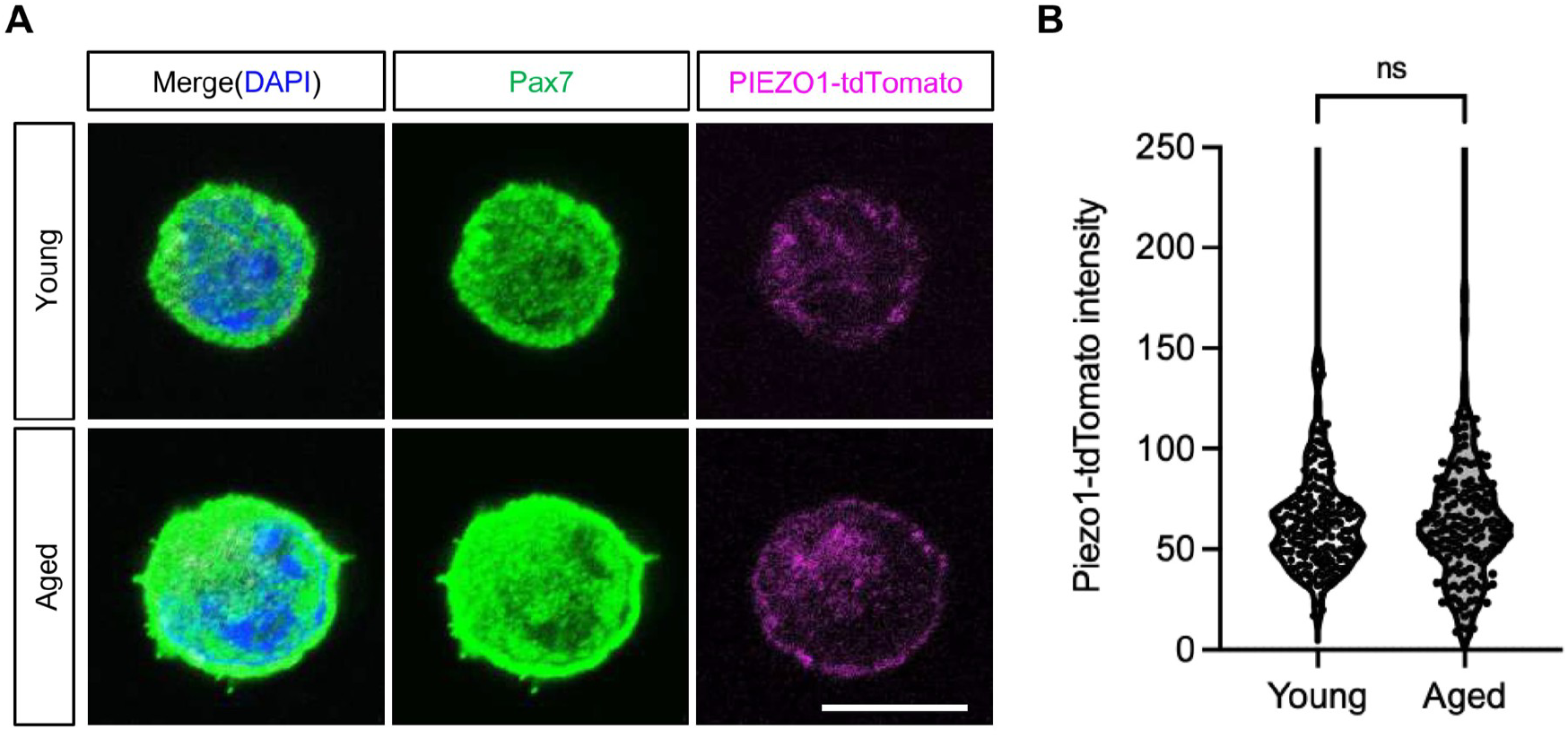
Expression of mechanosensitive ion channel PIEZO1 in MuSCs during muscle regeneration. **(A)** Representative immunofluorescence images of MuSCs isolated from young and aged PIEZO1-tdTomato mice at 3 days after CTX-induced muscle injury. Cells were stained for Pax7 and DAPI, and PIEZO1-tdTomato fluorescence was detected directly. Scale bar, 10 μm. **(B)** Quantification of PIEZO1-tdTomato fluorescence intensity in young and aged MuSCs. No significant difference was observed between young and aged MuSCs. Each dot represents an individual MuSC. Data are presented as violin plots with individual data points. n.s., not significant. Unpaired *t*-test.

## References

1. Ebashi, S. & Endo, M. Calcium ion and muscle contraction. Prog Biophys Mol Biol 18, 123–183, doi:10.1016/0079-6107(68)90023-0 (1968).

2. Bagur, R. & Hajnoczky, G. Intracellular Ca2+ Sensing: Its Role in Calcium Homeostasis and Signaling. Mol Cell 66, 780–788, doi:10.1016/j.molcel.2017.05.028 (2017)

3. Berridge, M. J., Lipp, P. & Bootman, M. D. The versatility and universality of calcium signalling. Nat Rev Mol Cell Biol 1, 11–21, doi:10.1038/35036035 (2000)

4. Berridge, M. J., Bootman, M. D. & Roderick, H. L. Calcium signalling: dynamics, homeostasis and remodelling. Nat Rev Mol Cell Biol 4, 517–529, doi:10.1038/nrm1155 (2003)

5. Ikegaya, Y., Le Bon-Jego, M. & Yuste, R. Large-scale imaging of cortical network activity with calcium indicators. Neurosci Res 52, 132–138, doi:10.1016/j.neures.2005.02.004 (2005)

6. Qian, N. et al. TRPM7 channels mediate spontaneous Ca2+ fluctuations in growth plate chondrocytes that promote bone development. Sci Signal 12, doi:10.1126/scisignal.aaw4847 (2019)

7. Liu, F., Xu, L., Nishi, M., Ichimura, A. & Takeshima, H. Enhanced Ca2+ handling in thioglycolate-elicited peritoneal macrophages. Cell Calcium 96, 102381, doi:10.1016/j.ceca.2021.102381 (2021)

8. Miyazaki, Y. et al. C-type natriuretic peptide facilitates autonomic Ca2+ entry in growth plate chondrocytes for stimulating bone growth. Elife 11, doi:10.7554/eLife.71931 (2022)

9. Arjun McKinney, A., Petrova, R. & Panagiotakos, G. Calcium and activity-dependent signaling in the developing cerebral cortex. Development 149, doi:10.1242/dev.198853 (2022)

10. Hirano, K. & Hara, Y. Mechanobiological landscape of muscle stem cells. Skelet Muscle, doi:10.1186/s13395-026-00425-6 (2026)

11. Gugliuzza, M. V. & Crist, C. Muscle stem cell adaptations to cellular and environmental stress. Skelet Muscle 12, 5, doi:10.1186/s13395-022-00289-6 (2022)

12. Dumont, N. A., Wang, Y. X. & Rudnicki, M. A. Intrinsic and extrinsic mechanisms regulating satellite cell function. Development 142, 1572–1581, doi:10.1242/dev.114223 (2015)

13. Fukada, S. et al. Purification and cell-surface marker characterization of quiescent satellite cells from murine skeletal muscle by a novel monoclonal antibody. Exp Cell Res 296, 245–255, doi:10.1016/j.yexcr.2004.02.018 (2004)

14. Liu, L., Cheung, T. H., Charville, G. W. & Rando, T. A. Isolation of skeletal muscle stem cells by fluorescence-activated cell sorting. Nat Protoc 10, 1612–1624, doi:10.1038/nprot.2015.110 (2015)

15. Feige, P. & Rudnicki, M. A. Isolation of satellite cells and transplantation into mice for lineage tracing in muscle. Nat Protoc 15, 1082–1097, doi:10.1038/s41596-019-0278-8 (2020)

16. Motohashi, N., Asakura, Y. & Asakura, A. Isolation, culture, and transplantation of muscle satellite cells. J Vis Exp, doi:10.3791/50846 (2014)

17. Machado, L. et al. In Situ Fixation Redefines Quiescence and Early Activation of Skeletal Muscle Stem Cells. Cell Rep 21, 1982–1993, doi:10.1016/j.celrep.2017.10.080 (2017)

18. van Velthoven, C. T. J., de Morree, A., Egner, I. M., Brett, J. O. & Rando, T. A. Transcriptional Profiling of Quiescent Muscle Stem Cells In Vivo. Cell Rep 21, 1994–2004, doi:10.1016/j.celrep.2017.10.037 (2017)

19. Machado, L. et al. Tissue damage induces a conserved stress response that initiates quiescent muscle stem cell activation. Cell Stem Cell 28, 1125–1135 e1127, doi:10.1016/j.stem.2021.01.017 (2021)

20. Kann, A. P. et al. An injury-responsive Rac-to-Rho GTPase switch drives activation of muscle stem cells through rapid cytoskeletal remodeling. Cell Stem Cell 29, 933–947 e936, doi:10.1016/j.stem.2022.04.016 (2022)

21. Cho, C. H., Woo, J. S., Perez, C. F. & Lee, E. H. A focus on extracellular Ca2+ entry into skeletal muscle. Exp Mol Med 49, e378, doi:10.1038/emm.2017.208 (2017)

22. Roman, W. et al. Muscle repair after physiological damage relies on nuclear migration for cellular reconstruction. Science 374, 355–359, doi:10.1126/science.abe5620 (2021)

23. Bansal, D., Miyake, K., Vogel, S. et al. Defective membrane repair in dysferlin-deficient muscular dystrophy. Nature 423, 168–172,doi: 10.1038/nature01573 (2003)

24. Tu, M. K., Levin, J. B., Hamilton, A. M. & Borodinsky, L. N. Calcium signaling in skeletal muscle development, maintenance and regeneration. Cell Calcium 59, 91–97, doi:10.1016/j.ceca.2016.02.005 (2016)

25. Eigler, T. et al. ERK1/2 inhibition promotes robust myotube growth via CaMKII activation resulting in myoblast-to-myotube fusion. Dev Cell 56, 3349–3363 e3346, doi:10.1016/j.devcel.2021.11.022 (2021)

26. Li, T. et al. STIM1-Ca2+ signaling is required for the hypertrophic growth of skeletal muscle in mice. Mol Cell Biol 32, 3009–3017, doi:10.1128/MCB.06599-11 (2012)

27. Ito, N., Ruegg, U. T., Kudo, A., Miyagoe-Suzuki, Y. & Takeda, S. Activation of calcium signaling through Trpv1 by nNOS and peroxynitrite as a key trigger of skeletal muscle hypertrophy. Nat Med 19, 101–106, doi:10.1038/nm.3019 (2013)

28. Franco, A., Jr. & Lansman, J. B. Calcium entry through stretch-inactivated ion channels in mdx myotubes. Nature 344, 670–673, doi:10.1038/344670a0 (1990)

29. Millay, D. P. et al. Calcium influx is sufficient to induce muscular dystrophy through a TRPC-dependent mechanism. Proc Natl Acad Sci U S A 106, 19023–19028, doi:10.1073/pnas.0906591106 (2009)

30. Rog, J., Oksiejuk, A., Gorecki, D. C. & Zablocki, K. Primary mouse myoblast metabotropic purinoceptor profiles and calcium signalling differ with their muscle origin and are altered in mdx dystrophinopathy. Sci Rep 13, 9333, doi:10.1038/s41598-023-36545-y (2023)

31. Hirano, K. et al. The mechanosensitive ion channel PIEZO1 promotes satellite cell function in muscle regeneration. Life Sci Alliance 6, doi:10.26508/lsa.202201783 (2023)

32. Ohkura, M. et al. Genetically encoded green fluorescent Ca2+ indicators with improved detectability for neuronal Ca2+ signals. PLoS One 7, e51286, doi:10.1371/journal.pone.0051286 (2012)

33. Murphy, M. M., Lawson, J. A., Mathew, S. J., Hutcheson, D. A. & Kardon, G. Satellite cells, connective tissue fibroblasts and their interactions are crucial for muscle regeneration. Development 138, 3625–3637, doi:10.1242/dev.064162 (2011)

34. Dos Santos, M. et al. A fast Myosin super enhancer dictates muscle fiber phenotype through competitive interactions with Myosin genes. Nat Commun 13, 1039, doi:10.1038/s41467-022-28666-1 (2022)

35. Ono, Y., Boldrin, L., Knopp, P., Morgan, J. E. & Zammit, P. S. Muscle satellite cells are a functionally heterogeneous population in both somite-derived and branchiomeric muscles. Dev Biol 337, 29–41, doi:10.1016/j.ydbio.2009.10.005 (2010)

36. Yoshioka, K. et al. Hoxa10 mediates positional memory to govern stem cell function in adult skeletal muscle. Sci Adv 7, doi:10.1126/sciadv.abd7924 (2021)

37. Motohashi, N. et al. Tbx1 regulates inherited metabolic and myogenic abilities of progenitor cells derived from slow- and fast-type muscle. Cell Death Differ 26, 1024–1036, doi:10.1038/s41418-018-0186-4 (2019)

38. Blau, H. M., Cosgrove, B. D. & Ho, A. T. The central role of muscle stem cells in regenerative failure with aging. Nat Med 21, 854–862, doi:10.1038/nm.3918 (2015)

39. Hirano, K. et al. Mg2+ influx mediated by TRPM7 triggers the initiation of muscle stem cell activation. Sci Adv 11, eadu0601, doi:10.1126/sciadv.adu0601 (2025)

40. Ma, N. et al. Piezo1 regulates the regenerative capacity of skeletal muscles via orchestration of stem cell morphological states. Sci Adv 8, eabn0485, doi:10.1126/sciadv.abn0485 (2022)

41. Konagaya, Y. et al. Intravital imaging reveals cell cycle-dependent myogenic cell migration during muscle regeneration. Cell Cycle 19, 3167–3181, doi:10.1080/15384101.2020.1838779 (2020)

42. Webster, M. T., Manor, U., Lippincott-Schwartz, J. & Fan, C. M. Intravital Imaging Reveals Ghost Fibers as Architectural Units Guiding Myogenic Progenitors during Regeneration. Cell Stem Cell 18, 243–252, doi:10.1016/j.stem.2015.11.005 (2016)

43. Siegel, A. L., Atchison, K., Fisher, K. E., Davis, G. E. & Cornelison, D. D. 3D timelapse analysis of muscle satellite cell motility. Stem Cells 27, 2527–2538, doi:10.1002/stem.178 (2009)

44. Choi, S., Ferrari, G. & Tedesco, F. S. Cellular dynamics of myogenic cell migration: molecular mechanisms and implications for skeletal muscle cell therapies. EMBO Mol Med 12, e12357, doi:10.15252/emmm.202012357 (2020)

45. SenGupta, S., Parent, C. A. & Bear, J. E. The principles of directed cell migration. Nat Rev Mol Cell Biol 22, 529–547, doi:10.1038/s41580-021-00366-6 (2021)

46. Ebashi, S. Calcium Binding and Relaxation in the Actomyosin System. J Biochem, doi:10.1093/oxfordjournals.jbchem.a127142 (1960).

47. Ebashi, S. Calcium binding activity of vesicular relaxing factor. J Biochem 50, 236–244, doi:10.1093/oxfordjournals.jbchem.a127439 (1961).

48. Ozawa, E. & Ebashi, S. Requirement of Ca ion for the stimulating effect of cyclic 3’,5’-AMP on muscle phosphorylase b kinase. J Biochem 62, 285–286, doi:10.1093/oxfordjournals.jbchem.a128663 (1967).

49. Ebashi, S., Ebashi, F. & Kodama, A. Troponin as the Ca++-receptive protein in the contractile system. J Biochem 62, 137–138, doi:10.1093/oxfordjournals.jbchem.a128628 (1967).

50. Colom, A. et al. A fluorescent membrane tension probe. Nat Chem 10, 1118–1125, doi:10.1038/s41557-018-0127-3 (2018)

51. Okabe, K. et al. Intracellular temperature mapping with a fluorescent polymeric thermometer and fluorescence lifetime imaging microscopy. Nat Commun 3, 705, doi:10.1038/ncomms1714 (2012)

52. Kiyonaka, S. et al. Genetically encoded fluorescent thermosensors visualize subcellular thermoregulation in living cells. Nat Methods 10, 1232–1238, doi:10.1038/nmeth.2690 (2013)

53. Miesenbock, G., De Angelis, D. A. & Rothman, J. E. Visualizing secretion and synaptic transmission with pH-sensitive green fluorescent proteins. Nature 394, 192–195, doi:10.1038/28190 (1998)

54. Zhang, Y. et al. Fast and sensitive GCaMP calcium indicators for imaging neural populations. Nature 615, 884–891, doi:10.1038/s41586-023-05828-9 (2023)

55. Rocheteau, P., Gayraud-Morel, B., Siegl-Cachedenier, I., Blasco, M. A. & Tajbakhsh, S. A subpopulation of adult skeletal muscle stem cells retains all template DNA strands after cell division. Cell 148, 112–125, doi:10.1016/j.cell.2011.11.049 (2012)

56. Carrillo-Garcia, J. et al. The mechanosensitive Piezo1 channel controls endosome trafficking for an efficient cytokinetic abscission. Sci Adv 7, eabi7785, doi:10.1126/sciadv.abi7785 (2021)

57. Lacraz, G. et al. Increased Stiffness in Aged Skeletal Muscle Impairs Muscle Progenitor Cell Proliferative Activity. PLoS One 10, e0136217, doi:10.1371/journal.pone.0136217 (2015).

58. Segel, M. et al. Niche stiffness underlies the ageing of central nervous system progenitor cells. Nature 573, 130–134, doi:10.1038/s41586-019-1484-9 (2019).

59. Wei, C. et al. Calcium flickers steer cell migration. Nature 457, 901–905, doi:10.1038/nature07577 (2009).

60. Shellard, A. & Mayor, R. Durotaxis: The Hard Path from In Vitro to In Vivo. Dev Cell 56, 227–239, doi:10.1016/j.devcel.2020.11.019 (2021).

61. Lu, H. H., Ege, D., Salehi, S. & Boccaccini, A. R. Ionic medicine: Exploiting metallic ions to stimulate skeletal muscle tissue regeneration. Acta Biomater 190, 1–23, doi:10.1016/j.actbio.2024.10.033 (2024)

62. Lepper, C., Conway, S. J. & Fan, C. M. Adult satellite cells and embryonic muscle progenitors have distinct genetic requirements. Nature 460, 627–631, doi:10.1038/nature08209 (2009)

63. Gorelik, R. & Gautreau, A. Quantitative and unbiased analysis of directional persistence in cell migration. Nat Protoc 9, 1931–1943, doi:10.1038/nprot.2014.131 (2014)

