## Supplementary figures and images for "Biomechanical regulation of Ca^2+^ dynamics during muscle stem cell activation"

### Supplemental Figure 1

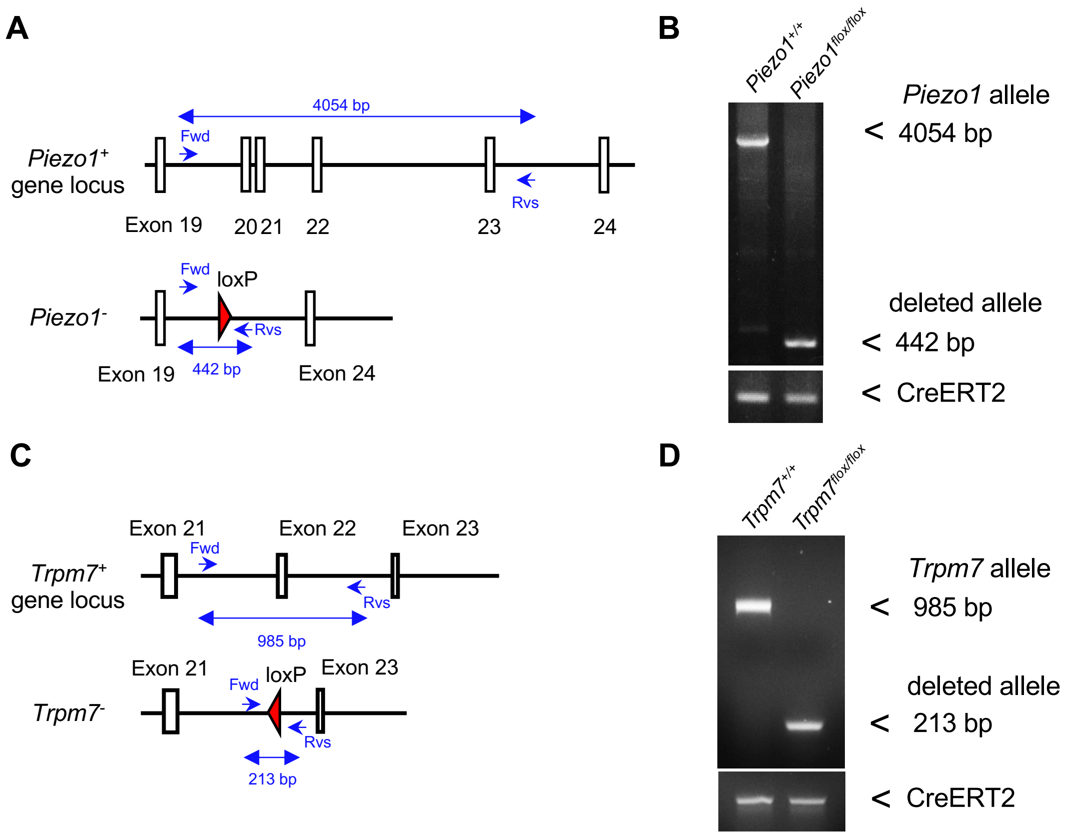

### Supplemental Figure 2

A

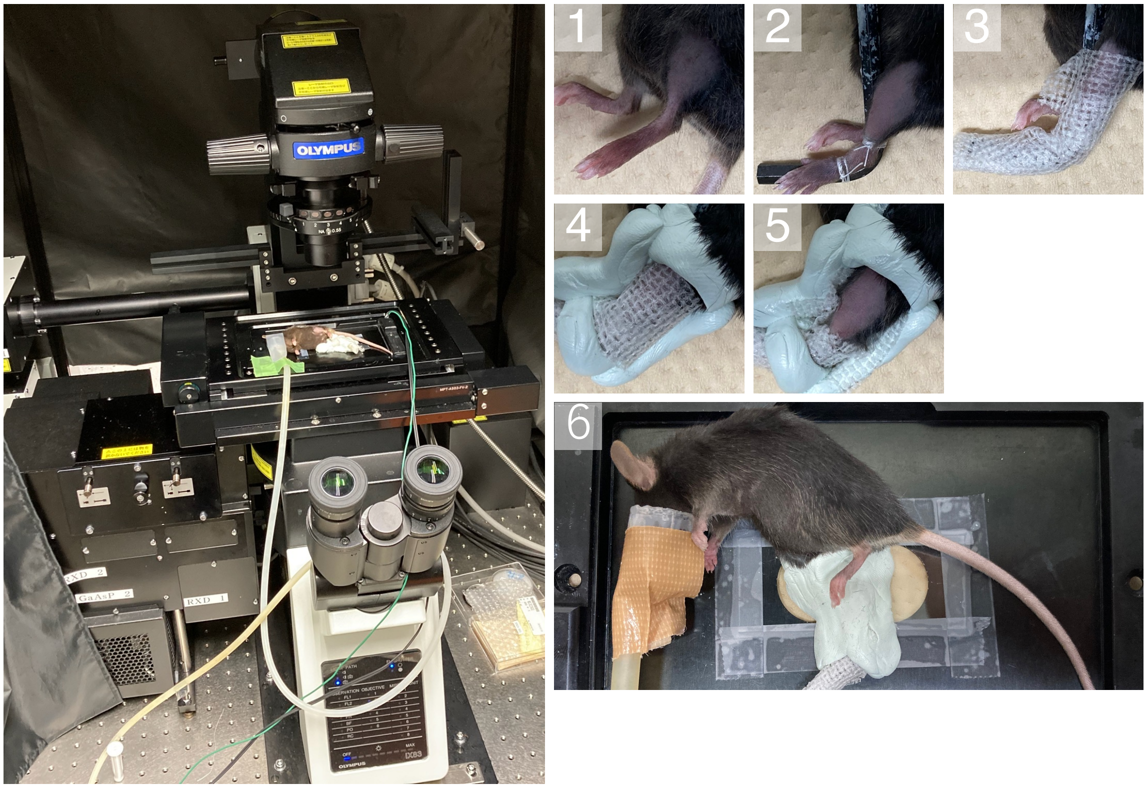


B

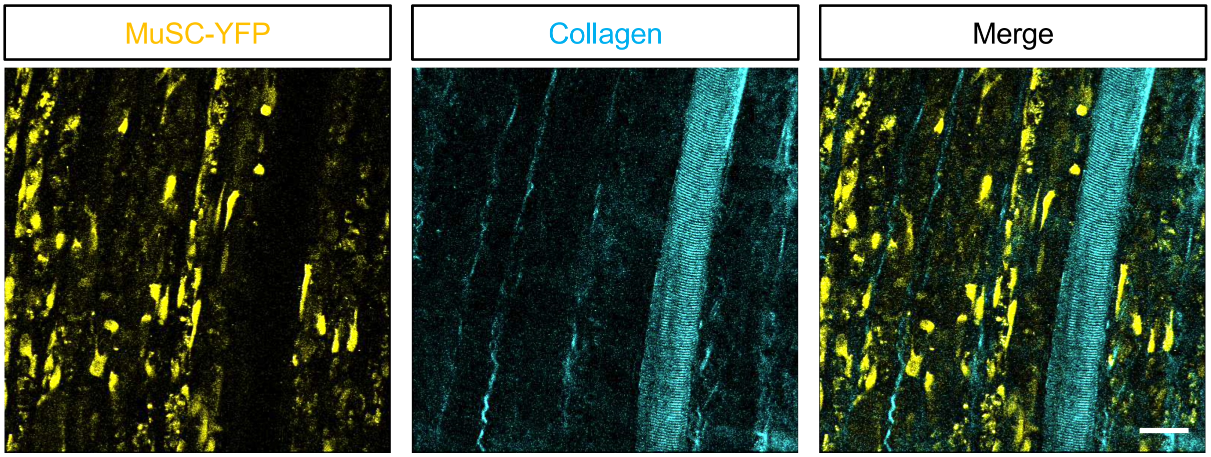

### Supplemental Figure 3

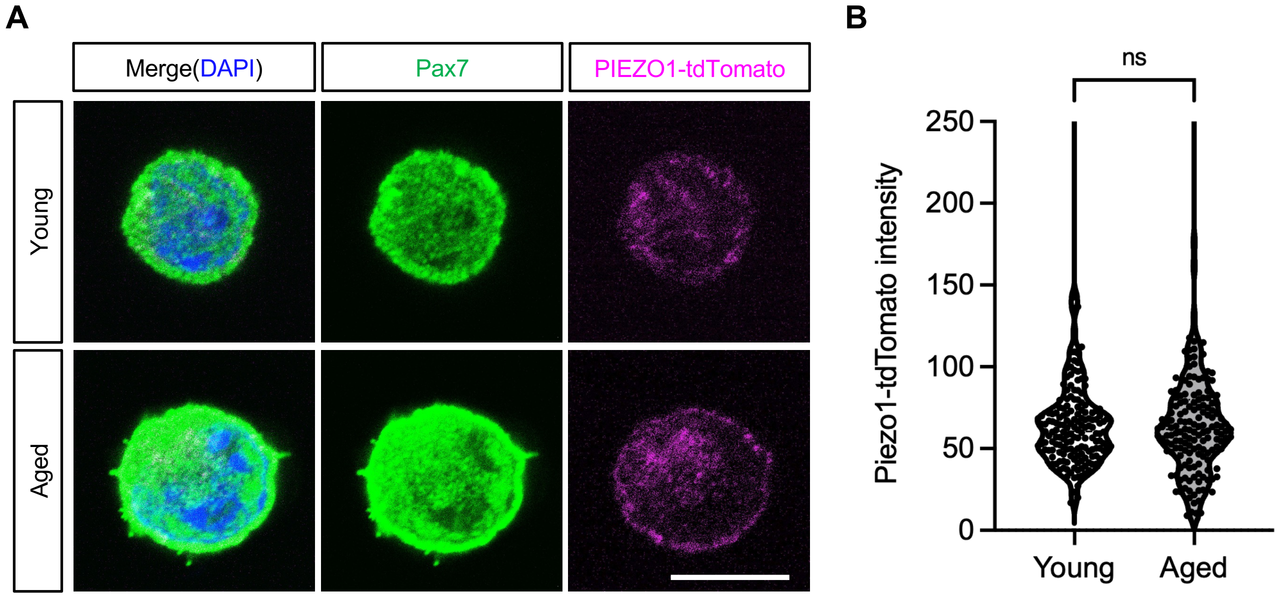
